# The mitogen-activated protein kinase kinase kinase, ILK5, regulates plant purinergic receptor-mediated, innate immunity

**DOI:** 10.1101/2022.04.19.488815

**Authors:** Daewon Kim, Dongqin Chen, Nagib Ahsan, Jay J. Thelen, Gary Stacey

**Affiliations:** Division of Plant Science and Technology, C.S. Bond Life Science Center, University of Missouri, Columbia, MO 65211 USA; Division of Biochemistry, C.S. Bond Life Science Center, University of Missouri, Columbia, MO 65211 USA; State Key Laboratory of Agrobiotechnology, College of Plant Protection, China Agricultural University, Beijing, 100193 China; Department of Chemistry and Biochemistry, The University of Oklahoma, Norman, OK 73019, USA; Mass Spectrometry, Proteomics and Metabolomics Core Facility, Stephenson Life Sciences Research Center, University of Oklahoma, Norman, OK, USA

**Keywords:** Mitogen-activated protein (MAP) kinase, Extracellular ATP, Integrin-linked kinase, ILK5, Purinergic signaling, P2K1, plant innate immunity

## Abstract

Mitogen-activated protein (MAP) kinase signaling cascades play important roles in the regulation of eukaryotic defense against various pathogens. Activation of the extracellular ATP (eATP) receptor P2K1 triggers MAP kinase 3 and 6 (MPK3/6) phosphorylation, which leads to elevated defense responses in Arabidopsis. However, the mechanism by which P2K1 activates the MAPK cascade is unclear. In this study, we identified Raf-like MAPKKK ILK5 as a downstream substrate of the P2K1 kinase. P2K1 phosphorylates ILK5 on serine 192. The interaction between P2K1 and ILK5 was confirmed both *in vitro* and *in planta* and their interaction was enhanced by ATP treatment. Similar to *P2K1* expression, *ILK5* expression levels were highly induced by treatment with ATP, flg22, *Pseudomonas syringae* pv. *tomato* DC3000, and various abiotic stresses, such as wounding. ILK5 interacts with both MKK4 and MKK5, but only MKK5 is phosphorylated by ILK5. Moreover, phosphorylation of MPK3/6 was significantly reduced upon ATP treatment in *ilk5* mutant plants, relative to wild-type. The *ilk5* mutant plants showed higher susceptibility to *P. syringae* pathogen infection relative to wild-type plants. Plants expressing only the mutant ILK5^S192A^ protein, lacking kinase activity, did not activate the MAPK cascade upon ATP addition. Taken together, the results suggest that eATP activation of P2K1 results in transphosphorylation of the Raf-like MAPKKK ILK5, which subsequently triggers the MAPK cascade, culminating in activation of MAPK3 and 6 associated with an elevated innate immunity response.

**Significance statement:** Pathogens invasion and subsequent wound stress activates extracellular ATP-mediated purinergic signaling cascades, a danger associated molecular pattern (DAMP) signal, which induces phosphorylation of MAPKs. Previous studies revealed that the P2K1 purinergic receptor increases MPK3/6 phosphorylation in response to eATP signaling cascades in Arabidopsis. However, the specific mechanism by which this occurs remains unknown. Here, we describe the isolation and characterization of Raf-like MAPKKK ILK5 (Integrin-linked Kinase 5) as a downstream substrate of P2K1 kinase activity. Initiation of an eATP-dependent signaling pathway by phosphorylation of ILK5 with subsequent activation of MKK5, leading to activation of MPK3/6 and downstream events is crucial to the plant innate immunity response.

## Introduction

Adenosine 5′-triphosphate (ATP) serves as an intracellular energy currency for all organisms. However, in both plants and animals, extracellular ATP (eATP) also functions as a danger-associated molecular pattern (DAMP) signal molecule when perceived by plasma-membrane localized receptors(1, 2). Mammals possess two major classes of purinoceptors, ligand-gated ion channel P2X and G protein-coupled receptor P2Y, which have been extensively studied(3). eATP has been shown to play a variety of roles in mammals, including regulating the immune response, inflammation, neurotransmission, muscle contraction, and cell death(4–6). In contrast to animals, the functional role of eATP in plants is not well studied. Nevertheless, published data suggest diverse roles for purinergic signaling in plants, including involvement in the response to biotic and abiotic stresses(7–10), gravitropism(11), root hair growth(12, 13), root avoidance(14), thigmotropism(15), and cell death(16, 17). Plants lack canonical P2X and P2Y receptors(18, 19). Thus, it was a significant breakthrough when the first plant purinergic receptor was identified as a member of the lectin receptor-like kinase family (i.e., LecRK I.9), originally termed DOESN’T RESPOND TO NUCLEOTIDES 1 (DORN1) but more recently named P2K1, in keeping with the nomenclature originally established in animals for P2-type purinoreceptors(8, 18). P2K1 is composed of an extracellular legume L-type lectin domain at the N-terminus, a trans-membrane domain at the middle region, and an intracellular serine/threonine kinase domain at the C-terminus(8, 20). P2K1 (LecRK I.9) was originally identified as a positive regulator of plant defense against the oomycete pathogens, *Phytophthora brassicae* and *Phytophthora infestans*, and the bacterial pathogen *Pseudomonas syringae* pv. *tomato* DC3000 (*Pst* DC3000) (8, 21–24). However, the primary biochemical function of P2K1 appears to be as a receptor during developmental and stress conditions that induce the release of eATP (e.g., upon wounding)(1, 19). Recently, it was demonstrated that P2K1 plays an important role in regulating the production of reactive oxygen species (ROS) via direct phosphorylation of NADPH oxidase (i.e., RBOHD) regulating both stomatal aperture and plant innate immunity(8). Indeed, very recently it was shown that eATP triggers a ROS wave in plants that is dependent on P2K1 function(25). It was also shown that S-acylation affects the temporal dynamics of P2K1 receptor activity through autophosphorylation and protein degradation. The CYCLIC NUCLEOTIDE GATED CHANNEL 6 (CNGC6) and CNGC2 proteins were shown to play a crucial role in mediating eATP-induced cytosolic Ca^2+^ signaling(26–28). Animals possess multiple P2X and P2Y receptors and, hence, it was perhaps no surprise when a second, plant purinoreceptor, P2K2 (LecRK I.5) was identified(29). Both P2K1 and P2K2 bind ATP and ADP with high affinity(18, 29). Among the various downstream events activated by eATP is phosphorylation of MAP kinase 3 and 6, which is markedly reduced in *p2k1/p2k2* mutant plants(18, 29).

The mitogen-activated protein kinase (MAPK) cascade plays a critical role in transmitting and amplifying stimulus-specific signals to the cellular machinery by phosphorylation of target proteins in eukaryotes(30, 31). Compared to the small number of plant MAPKK and MAPK family members, MAPKKK constitute a larger family, which are considered essential to the ability to respond to diverse signals/environmental conditions(31, 32). For example, the Arabidopsis genome encodes genes for 20 MAPK, 10 MAPKK and 80 MAPKKK subfamily members(31, 32). Plant MAPKKK are divided into three subfamilies, MEKK, Raf and ZIK. In particular, the plant Raf-like kinase (Raf) family is much larger in number and more structurally diverse compared to metazoans(33). Among 80 MAPKKKs in Arabidopsis, 21, 48 and 11 members belong to MEKK, Raf, and ZIK subfamilies, respectively(34). However, the biological function of only a small fraction of the Raf-like MAPKKK is known. Among the Raf-like MAPKKK members, Enhanced Disease Resistance 1 (EDR1/Raf2) and Constitutive Triple Response 1 (CTR1/Raf1) are relatively well characterized(35, 36). CTR1 encodes a serine/threonine kinase that negatively regulates ethylene signaling by acting upstream of MKK9 and MPK3/6(37–39). In addition, mutational loss of CTR1 function causes defects in the sugar response(40). Another Raf-like MAPKKK, EDR1 also acts as a negative regulator in ethylene responses similarly to CTR1(36, 41). EDR1 regulates MAPK cascades via direct association with MKK4/MKK5 for negative regulation of salicylic acid (SA)-inducible, plant defense responses(36). *edr1* mutant plants display enhanced cell death in response to various abiotic stress conditions, such as drought and aging(36, 41). Integrin-linked Kinases (ILKs), a Raf MAPKKK subfamily with unique characteristics, are thought to play an important role in diverse biological functions in plants(42). The first ILK described was shown to be an interacting partner of the integrin receptor cytoplasmic *β* subunit, functioning in the assembly of signaling complexes at plasma membrane focal adhesion regions in metazoans(43, 44). ILKs were originally characterized as serine/threonine kinases, but recent studies have questioned whether ILKs actually function as kinases(45, 46). Instead, ILKs were proposed to function as scaffolding proteins with adaptor proteins, PINCH and α-parvin (IPP complex), to regulate various biological processes, such as cell adhesion, migration, proliferation, differentiation, assembly of the extracellular matrix (ECM), contractility, etc(43–47). In contrast to metazoans, plant genomes encode multiple ILK genes [e.g., 6 members, ILK1-6, in Arabidopsis](42). Plant ILKs were first described as N-terminus ankyrin repeat containing kinases suggested to play a role in osmotic stress and adventitious root growth in *Medicago* and *Arabidopsis*(48, 49). Previous studies showed that the vascular specific adaptor protein ILK6/VH1-INTERACTING KINASE (VIK) is involved in auxin and brassinosteroid signaling to regulate leaf venation(50). Recently, ILK1 was demonstrated to modulate the sensitivity of plants to osmotic and saline stress, response to nutrient availability, and response to bacterial pathogens(42, 51). While metazoan ILKs have been studied intensively, their role in molecular signaling in plants is poorly understood(42).

In this study, we identified Raf-like MAPKKK ILK5/Raf27/BHP (hereafter referred to as ILK5), previously characterized as a regulator of blue light-dependent stomatal movement(52), as a protein substrate of the P2K1 kinase. Indeed, P2K1 directly phosphorylates ILK5 on Ser192. Interaction between P2K1 and ILK5 was confirmed using a variety of *in vitro* and *in planta* methods. Taken together, we reveal that the presence of an eATP-dependent signaling pathway initiated by phosphorylation of ILK5 with subsequent activation of MKK5, leading to activation of MPK3/6 and downstream events crucial to the plant innate immunity response.

## Results

### ILK5 was identified as a P2K1 substrate

In order to identify substrate proteins for the P2K1 kinase domain, a mass spectrometry-based *in vitro* phosphorylation strategy, termed kinase client assay (KiC assay)(53, 54), was used. A synthetic peptide library, representing approximately 2100 experimentally identified *in vivo* phosphorylation sites, was incubated with GST-fused P2K1 intracellular domain recombinant protein in the presence of ATP. Subsequently, tandem mass spectrometry was performed to identify both phospho-peptides and specific phosphorylation sites. Two empty vectors (GST and MBP) and two P2K1 inactive kinase proteins [GST fused P2K1-KD-1 (D572N) and P2K1-KD-2 (D525N)] were used as negative controls. The result of this experiment was the identification of twenty-three phosphorylated peptides, which were identified and verified by phosphoRS score and phosphoRS site probability. As previously published, three protein substrates identified from this screen were RBOHD and PAT5/9, which were subsequently shown to be regulated by P2K1 phosphorylation(8, 26). In addition, the VKKLDDEVLS(p) peptide of ILK5 was phosphorylated by the P2K1 kinase domain (*Fig. 1A*). According to the Arabidopsis genome annotation database (https://www.arabidopsis.org/portals/genAnnotation/), BLAST search (www.ncbi.nlm.nih.gov./Blast/) and Pfam database (https://pfam.xfam.org/), *ILK5* encodes a Raf-like MAP kinase kinase kinase (MAPKKK) protein with a length of 459 amino acids and a molecular mass of approximately 50 kDa. ILK5 has the typical Raf-like MAPKKK structure with a N-terminal ankyrin repeat domain and C-terminal Ser/Thr kinase domain. ILK5/BHP was previously characterized as a regulator of blue light-dependent stomatal movement(52).

**Fig. 1.**
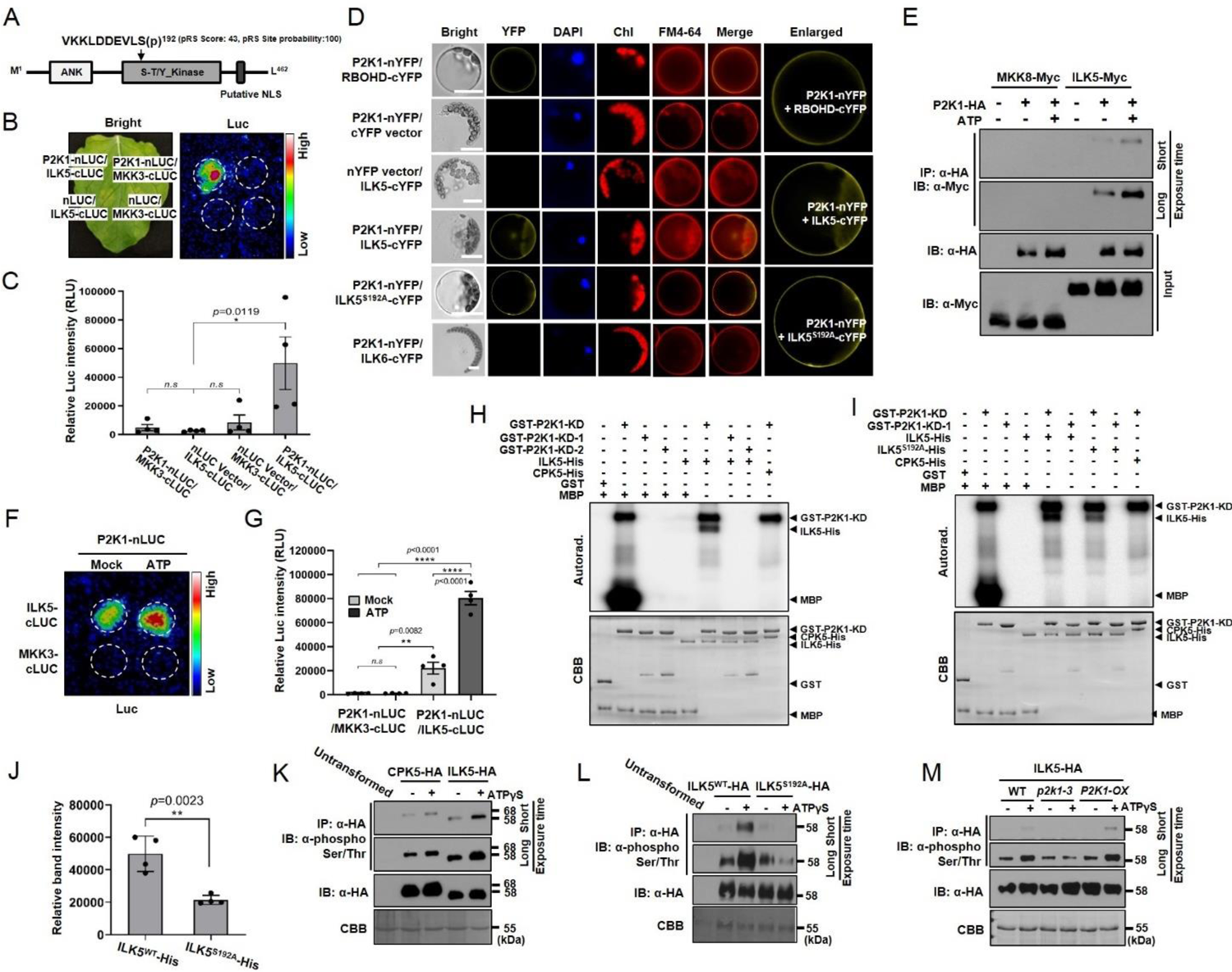
P2K1 interacts with and directly phosphorylates ILK5. (*A*) Schematic diagram of ILK5 (At4g18950) showing the ankyrin repeat (ANK) and serine-threonine or tyrosine kinase (S-T/Y kinase) domains. Identified phosphopeptide VKKLDDEVLS(p) is located in the kinase domain. (*B, C*, *F, G*) LCI experiment showing interaction of P2K1-ILK5 protein with/without 200 μM ATP treatment. The luminescence was monitored and captured using a low light imaging CCD camera (Photek; Photek, Ltd.). Dotted circles indicate the infiltrated area in *N. benthamiana* leaves. MKK3 protein was used as a negative control. (*D*) BiFC assay in Arabidopsis protoplasts. The YFP fluorescence was monitored using a Leica DM 5500B Compound Microscope with Leica DFC290 Color Digital Camera 24 h after transformation. DAPI and FM464 were used as a nuclear marker and plasma membrane marker, respectively. Chl represents chlorophyll auto-fluorescence signal. Merge indicates overlapped image of YFP and FM4-64. RBOHD and ILK6 were used as positive and negative control, respectively. Enlarged indicates enlargement of YFP image to clearly indicate plasma membrane-associated yellow fluorescence. Scale bars: 10 μm. (*E*) Co-immunoprecipitation of P2K1 and ILK5 proteins. The indicated constructs were co-infiltrated and transiently expressed in *N. benthamiana* leaves after addition of 200 μM ATP (+) for 30 min or MES buffer (pH 5.7) as a mock treatment (−). Total protein was used for Co-IP. Anti-HA and anti-Myc antibodies were used. MKK8-Myc was used as a negative control. (*H, I*) P2K1 directly phosphorylates ILK5 at Ser192. Purified recombinant ILK5-His and ILK5^S192A^ protein was incubated with GST-P2K1 kinase domain (GST-P2K1-KD), P2K1 kinase dead versions, (GST-P2K1-KD-1; D572N or GST-P2K1-KD-2; D525N), or GST in an *in vitro* kinase assay. Auto- and trans-phosphorylation were detected by incorporation of γ-[^32^P]-ATP. MBP and CPK5 were used as a universal substrate and a negative control, respectively. Protein loading was visualized by coomassie brilliant blue (CBB) staining. (*J*) Quantification of phosphorylated ILK5 and ILK5^S192A^ protein. The intensity of the phosphorylation signals of ILK5 and ILK5^S192A^ by P2K1-KD (shown in *SI Appendix, Fig. S2B*) were measured and analyzed using the Image J and GraphPad Prism 8. Data shown as mean ± SD, n = 4, \*\**p*<0.01, unpaired Student’s *t* test, two-way. (*K*) ILK5 protein is phosphorylated by ATP treatment *in vivo*. CPK5-HA used as a negative control. (*L*) Mutation of ILK5 at Ser192 results reduced phosphorylation under ATPγS treatment. (*M*) ILK5 phosphorylation is dependent on P2K1 protein. In panel *K*-*M*, either ILK5-HA or ILK5^S192A^-HA protein was expressed in Arabidopsis WT, *p2k1-3* or *P2K1-OX* protoplasts after addition of 250 μM ATPγS and subsequently immunoprecipitated (IP) using anti-HA antibody beads and immunoblotting (IB) was carried out with an anti-phospho-Ser/Thr antibody. Protein loading was visualized by CBB staining. Above experiments were repeated at least two times with similar results.

### P2K1 interacts with and directly phosphorylates ILK5

Phosphorylation of ILK5 by P2K1 in the KiC assay is consistent with direct interaction between these two proteins. To provide evidence for this interaction, we performed split-luciferase complementation imaging (LCI) in *Nicotiana benthamiana* leaves (*Fig. 1B* and *C*), with MKK3 serving as a negative control. In order to further confirm the interaction of P2K1 with ILK5 at the plasma membrane, bimolecular fluorescence complementation (BiFC) assays were conducted in Arabidopsis protoplasts. In order to confirm the subcellular localization of ILK5 in Arabidopsis, a YFP tagged ILK5 (C-term or N-term) was expressed in Arabidopsis transgenic plants and protoplasts. The bulk of ILK5 was detected in the cytosol but some fraction also co-localized with the FM4-64 and DAPI stains, localized to the plasma membrane and nucleus, respectively (*SI Appendix,* Fig. S1). In BiFC analysis, co-expression of P2K1-nYFP with ILK5-cYFP produced a yellow fluorescent signal that co-localized with the plasma membrane marker FM4-64, whereas co-expression of P2K1-nYFP with the control ILK6-cYFP protein did not display a yellow fluorescent signal (*Fig. 1D*). In these experiments, P2K1 interaction with RBOHD was used as a positive control (8). Hence, the data are consistent with a direct interaction of P2K1 and ILK5 at the plasma membrane. To provide further evidence for this interaction, we co-expressed HA-tagged P2K1 with a Myc-tagged ILK5 in *N. benthamiana* leaves, followed by immunoprecipitation using antibodies to the specific epitope tags. These experiments were done in the presence and absence of exogenous eATP. Consistent with a direct interaction, P2K1 and ILK5 were co-immunoprecipitated and their interaction appeared to be enhanced in the presence of eATP (*Fig. 1E*). We further confirmed these results using LCI in *N. benthamiana* leaves (*Fig. 1F* and *G*). It was previously reported that P2K1 has strong kinase activity, which is essential for eATP-induced, MPK3/6 phosphorylation *in planta*(8, 18). In order to confirm the results of the KiC assay, purified recombinant ILK5 protein was incubated with either purified GST-P2K1-KD, GST-P2K1-KD-1 or GST-P2K1-KD-2 in the presence of [γ-^32^P] ATP. The results of this assay showed that GST-P2K1-KD strongly trans-phosphorylated ILK5-His and the generic substrate MBP, but not CPK5 used as a negative control. Assays performed with the two kinase-dead versions of GST-P2K1-KD-1 and GST-P2K1-KD-2 failed to phosphorylate ILK5-His (*Fig. 1H*). To verify that the radiography signal was the result of phosphorylation, the addition of lambda protein phosphatase (Lambda PPase) was shown to reduce both auto- and trans-phosphorylation of P2K1 and ILK5 (*SI Appendix, Fig. S2A*). The tandem MS results from the KiC assay revealed Ser192 of ILK5 was the specific target of P2K1 phosphorylation. To confirm this, site-directed mutagenesis was used to generate an ILK5^S192A^ recombinant protein, which was purified and incubated with purified GST-P2K1-KD. As expected, the results showed a significant reduction in phosphorylation of ILK5^S192A^ recombinant protein (*Fig. 1I* and *J* and *SI Appendix, Fig. S2B*), relative to the wild-type protein. To investigate phosphorylation of ILK5 *in vivo*, ILK5-HA protein was expressed in Arabidopsis protoplasts and subsequently immunoprecipitated with an anti-HA antibody bead then probed by western blotting with an anti-phospho-Ser/Thr antibody. ILK5 phosphorylation was detected strongly under ATPγS (poorly hydrolyzed ATP analog) treatment. (*Fig. 1K*). In addition, mutation of ILK5 at Ser192 (ILK5^S192A^) led to significantly reduced phosphorylation by P2K1 *in vivo* (*Fig. 1L*). To investigate whether ILK5 phosphorylation is dependent on P2K1, ILK5-HA protein was expressed in wild-type, *p2k1-3* mutant and *P2K1* overexpressing transgenic plants under ATPγS treatment followed by immunoprecipitation using anti-HA antibody. Western blotting was subsequently performed with an anti phospho-Ser/Thr antibody. Reduction of phosphorylation and increased phosphorylation in ILK5-HA protein were detected in *p2k1-3* and *P2K1*-*OX* compared to wild-type respectively (*Fig. 1M*). In order to test whether this phosphorylation affected P2K1-ILK5 interaction, we conducted the LCI assay by co-expressing wild-type P2K1 protein with the ILK5 phospho-null (ILK5^S192A^) and phospho-mimic (ILK5^S192D^) protein forms (*SI Appendix,* Fig. S3). The results of this experiment suggest that the P2K1-ILK5 interaction is not dependent on the phosphorylation status of the ILK5 Ser192.

### ILK5 has kinase activity and activates MKK5

ILKs are involved in various processes in many organisms, however, kinase activity is not essential to the regulation of these processes(45, 55). Indeed, there is some dispute whether animal ILKs possess kinase activity(45). However, in contrast to the animal situation, an examination of the plant ILK protein sequences suggest that the core residues necessary for kinase activity are well conserved(42). These results suggest that ILK5, found in the Group C cluster of the Raf-like subfamily, may possess kinase activity(42). However, a previous study failed to find ILK5 kinase activity using protein purified after expression in bacterial cells(52). Consistent with the previous study(52), ILK5 protein purified from *E. coli* showed no or very weak kinase activity when assayed under a variety of conditions (data not shown). However, in contrast to protein expressed in bacteria, a previous study showed that ILK1 protein extracted from plant tissue had strong kinase activity(51). These results are also consistent with the report that ILK5 protein expressed and purified from wheat germ cells possessed strong kinase activity(56). Therefore, we expressed and purified the His-tagged ILK5 protein from *N. benthamiana* leaves. The ILK5 protein purified in this way was able to auto- and trans-phosphorylate myelin basic protein (MBP), a widely used universal substrate for kinase activity assays (*SI Appendix, Fig. S4A, B*). To determine if phosphorylation of ILK5 at S192 is required for kinase activity, *in vitro* kinase assays using phosphor-null (S192A) and phosphor-mimic (S192D) mutants were carried out. It was observed that S192A version of the ILK5 dramatically reduced kinase activity while the S192D mutation slightly increased kinase activity compared to WT ILK5 protein (*SI Appendix, Fig. S4C*). P2K1 autophosphorylation was used as a positive control. Taken together, we can conclude that ILK5 is an active kinase.

If ILK5 acts as a MAPK kinase kinase, one would expect it to interact with and activate downstream MAPK kinases. As a means to analyze this possibility, we first performed LCI analyses of ILK5 in the presence of each of the ten Arabidopsis MAPKKs (MKK1-10). Among these ten MKKs tested, ILK5 appeared to show strong interactions with both MKK4 and MKK5 (*Fig. 2A* and *B* and *SI Appendix, Fig. S5A*). MKK4 and MKK5 were previously shown to regulate phosphorylation of MPK3/6 and, hence, play an important role in regulating the plant immune response(57, 58). We further confirmed the interaction of ILK5 with both MKK4 and MKK5 by BiFC analysis. YFP signals were observed mainly in the cytoplasm and some nuclei of plants carrying either ILK5-cYFP and MKK4-nYFP or ILK5-cYFP and MKK5-nYFP, indicating that interactions between ILK5 and MKK4 or MKK5 took place in the cytoplasm and nucleus (*Fig. 2C*). To investigate further this interaction under ATP treatment, we co-expressed MKK4- and MKK5-nLUC with ILK5-cLUC and monitored luciferase intensity in the presence and absence of exogenous eATP. Leaves treated with ATP showed slightly enhanced MKK4-ILK5 protein interaction, but markedly increased MKK5-ILK5 protein interaction (Fig. 2 *D* and *E*). Furthermore, in order to examine whether ILK5 directly phosphorylates MKK4 or MKK5, *in vitro* kinase assays were performed with His-tagged ILK5 extracted from *N. benthamiana* plants and GST-tagged MKK4 and MKK5. In previous reports, MKK4 and MKK5 were shown to have strong auto- and trans-phosphorylation activities(59). Therefore, kinase dead versions of MKK4 (MKK4^K108R^) and MKK5 (MKK5^K99R^) were generated by site-directed mutagenesis according to a previous study(59) (*SI Appendix, Fig. S5B*). *In vitro* kinase assays were performed with ILK5 protein and kinase dead versions of MKK4 (MKK4^K108R^) or MKK5 (MKK5^K99R^). The results clearly showed that ILK5 directly phosphorylates MKK5^K99R^, while phosphorylation of MKK4^K108R^ was not detectable. (*Fig. 2F*). Previously, it was shown that MKK5 phosphorylation of Thr215 and Ser221 in the activation loop is required for MPK3/6 activation(59). A GST-MKK5^K99RT215AS221A^ was generated using site-directed mutagenesis. Reduced phosphorylation of GST-MKK5^K99RT215AS221A^ was detected compared with GST-MKK5^K99R^ (*Fig. 2G*). When Ser192 of ILK5 was substituted with Ala (ILK5^S192A^), phosphorylation of ILK5 by P2K1 was significantly reduced (*Fig. 1I* and *J*). Phosphorylation of MKK5^K99R^ protein was significantly reduced when incubated with the ILK5^S192A^ protein (*Fig. 2H*). To further investigate whether MKK5 phosphorylation is dependent on the ILK5 protein, MKK5-HA protein was expressed in wild-type and *ilk5-1* mutant plants under ATP treatment followed by immunoprecipitation using an anti-HA antibody. Western blotting was subsequently performed with an anti-phospho-Ser/Thr antibody. Reduced phosphorylation in MKK5-HA was detected in *ilk5-1* compared to wild-type in the presence of exogenous ATP (*Fig. 2I*). To verify how this phosphorylation affects their interaction, a LCI assay was performed between ILK5^S192A^ or ILK5^S192D^ and MKK4 or MKK5. These results indicate that the phosphorylation status of ILK5 does not significantly affect its ability to interact with MKK4 or MKK5 (*SI Appendix, Fig. S5C* and *D*).

**Fig. 2.**
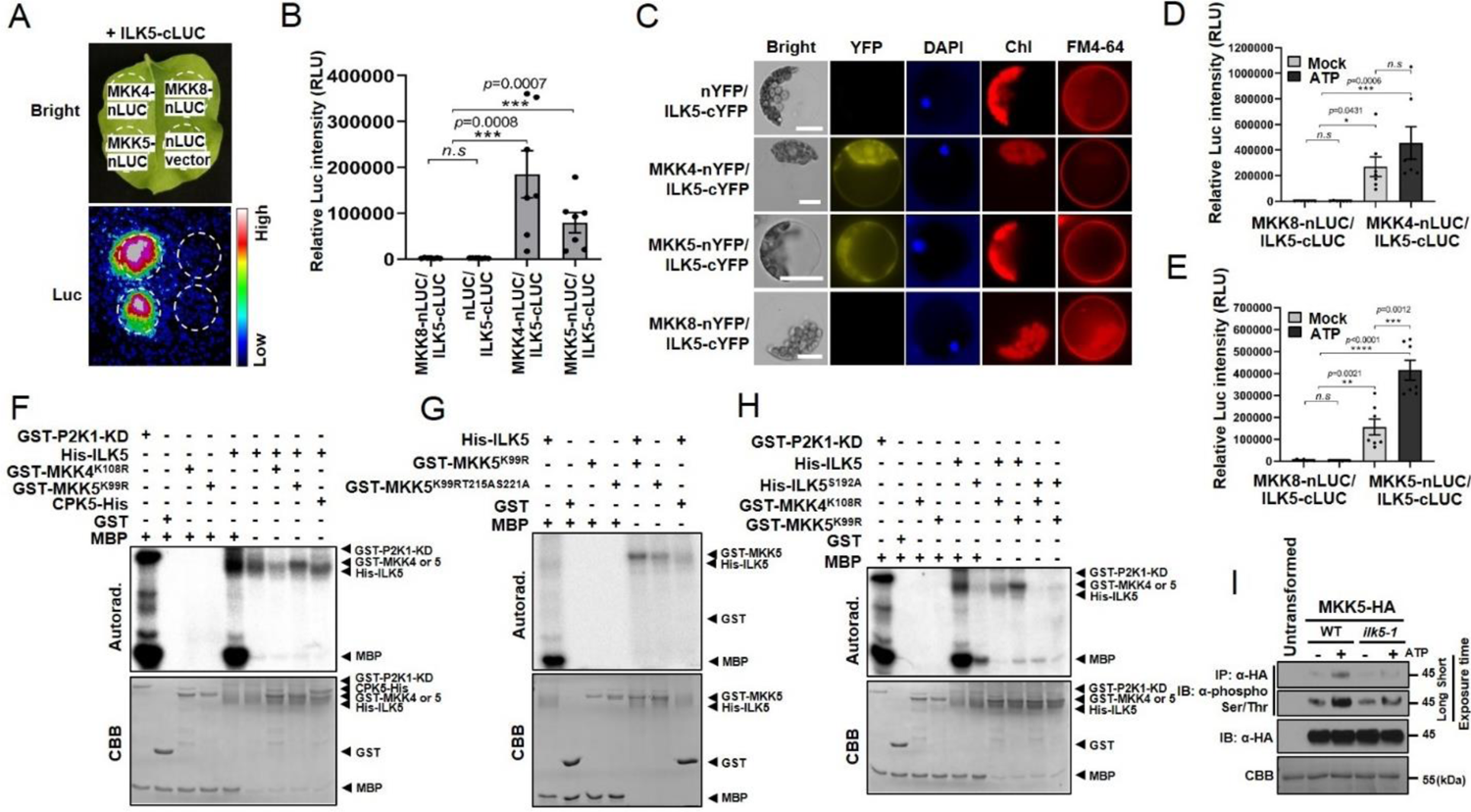
ILK5 interacts with MKK4 and MKK5 proteins. (*A*) ILK5 interaction between MKK4 and MKK5 was demonstrated using LCI analysis. Dotted circles indicate the infiltrated area in *N. benthamiana* leaves. MKK8 was used as a negative control. (*B*) Quantification of ILK5-MKK4 and -MKK5 interaction signal intensities. (*C*) BiFC assay in Arabidopsis protoplasts demonstrating ILK5 interaction with MKK4 and MKK5. DAPI and FM4-64 were used as nuclear and plasma membrane markers, respectively. Chl represents chlorophyll auto-fluorescence signal. MKK8 was used as a negative control. Scale bars: 10 μm. (*D* and *E*) Demonstration of ILK5-MKK4 and -MKK5 interaction signal intensities under ATP treatment. ILK5-MKK4 and -MKK5 interaction was monitored, images were captured, and luciferase signal intensities were quantified using C-vision/Im32 and analyzed using the GraphPad Prism 8. Data shown as mean ± SEM, n = 7 (biological replicates), **** *p*<0.0001, *** *p*<0.001, ** *p*<0.01, one-way ANOVA followed by Dunnett’s multiple comparisons. (*F-H*) MKK5 activation loop can be phosphorylated by ILK5. Recombinant His-ILK5 or His-ILK5^S192A^ protein purified from *N. benthamiana* was incubated with kinase dead GST-MKK4 (GST-MKK4^K108R^), kinase dead GST-MKK5 (GST-MKK5^K99R^), GST-MKK5 triple mutation (GST-MKK5^K99RT215AS221A^), CPK5-His or GST in an *in vitro* kinase assay. Autophosphorylation and trans-phosphorylation were detected by incorporation of γ-[^32^P]-ATP. MBP and CPK5 were used as an universal substrate and negative control, respectively. The protein loading was visualized by CBB staining. (*I*) MKK5 phosphorylation is reduced in *ilk5-1* mutant plants. MKK5-HA protein was expressed in wild-type and *ilk5-1* protoplasts with/without 250 μM ATP treatment, then subjected to IP and IB using anti-HA and anti-phospho-Ser/Thr antibodies. Protein loading was visualized by CBB staining. All above experiments were repeated at least two times with similar results.

### *ILK5* expression is strongly induced by ATP, wounding and pathogen treatments

In order to investigate the expression patterns of *ILK5* under comparable stress conditions, we stably transformed either a *ILK5promoter::GUS* or *ILK5promoter::GFP* construct into Arabidopsis. A published study indicated that *ILK5* is highly expressed in guard cells(52). Consistent with this published work, we also found strong *ILK5promoter::GUS* and *GFP* expression in guard cells (*Fig. 3A* and *SI Appendix, Fig. S6A*). The GUS signals observed in *ILK5promoter::GUS* transgenic plants showed ubiquitous expression in various tissues including young rosette leaves, primary root and shoot apices under normal growth conditions (*SI Appendix, Fig. S6B*). In order to directly compare to expression of the *P2K1* gene, GUS expression in *P2K1promoter::GUS* transgenic plants was also examined(60). This experiment showed that the pattern of *P2K1* expression was very similar, if not identical, to that found for *ILK5* expression under normal growth conditions (*SI Appendix, Fig. S6B* and *C*). Subsequently, qRT-PCR was use to confirm expression of both *ILK5* and *P2K1* expression in various tissues and abiotic stress conditions (*SI Appendix, Fig. S6D* and *E*).

**Fig. 3.**
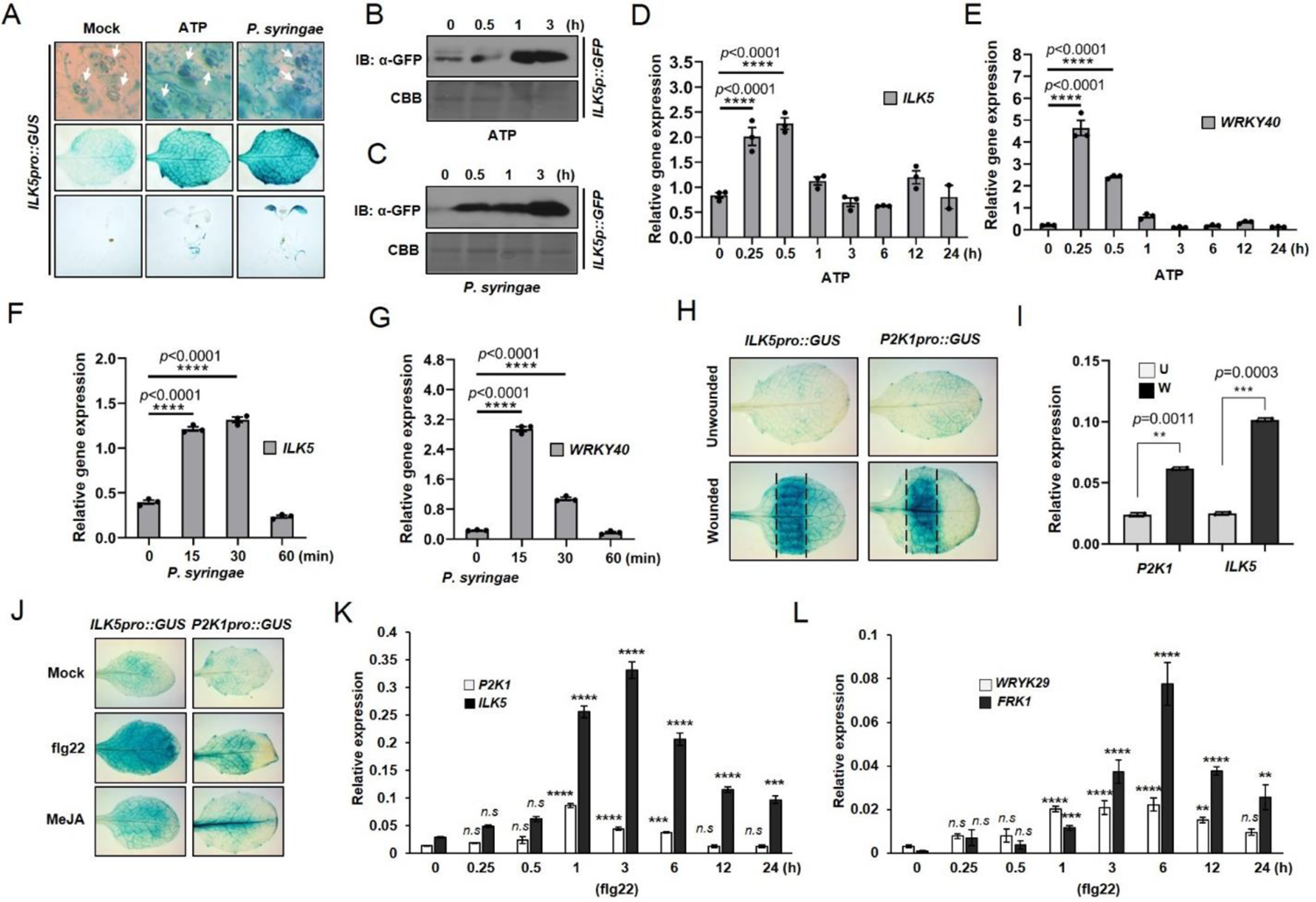
*ILK5* is highly induced after addition of ATP or *Pst* DC3000. (*A*) Histochemical analysis of *ILK5* expression after ATP treatment or *Pst* DC3000 inoculation. The expression patterns of the *ILK5promoter::GUS* transgenic plant was detected by histochemical staining in a 10-day-old seedling rosette leaf treated with 200 μM ATP or inoculated with *Pst* DC3000 (OD_600_ = 0.05) after 1 hour. Arrows indicate the stained guard cells. (*B*, *C*) Western blot analysis of GFP expressed from the *ILK5* promoter after ATP treatment or *Pst* DC3000 inoculation. Total protein was extracted from 10-day-old seedlings at each time point after treatments. Western blotting was performed using an anti-GFP antibody. CBB used as a loading control. (*D*-*G*) Real-time qRT-PCR analysis of *ILK5* transcripts after ATP (200 μM) or *Pst* DC3000 treatment. Total RNA was isolated from 10-days old seeding plants at each time point and 2 μg of total RNAs were used in this experiment. *WRKY40* was used as an inducible marker gene for the response to ATP and *Pst* DC3000 treatment in *Fig. 3E* and *G,* respectively. The *SAND* reference gene was used for data normalization. (*H*) Histochemical analysis of *ILK5promoter::GUS* and *P2K1promoter::GUS* expression in response to wounding. The expression patterns of the *ILK5promoter::GUS* and *P2K1promoter::GUS* transgenic plants were detected by histochemical staining of 2-week-old GUS transgenic plants after wounding. (*I*) Real-time qRT-PCR analysis of *ILK5* and *P2K1* gene expression in response to wounding. Data shown as mean ± SEM, ***P<0.001, **P<0.01, unpaired Student’s *t* test, two-way. (*J*) Histochemical analysis of 10-day-old seedling *ILK5promoter::GUS* and *P2K1promoter::GUS* in response to flg22(100 nM) and MeJA(1 μM). (*K*, *L*) Real-time qRT-PCR analysis of *ILK5* and *P2K1* gene expression (n = 3) at different time points in response to flg22 treatment. *FRK1* and *WRKY29* were used as inducible marker genes. In panel D, E, F, G, K and L, data shown as mean ± SEM, \*\*\*\**p*<0.0001, \*\*\**p*<0.001, \*\**p*<0.01, \**p*<0.05, *p*-value was determined and analyzed using the GraphPad Prism 8 by one-way ANOVA followed by Dunnett’s multiple comparisons. These experiments were repeated three times with similar results.

Previous published work clearly showed an important role for P2K1 in mediating the plant defense response to various pathogens, including wounding and jasmonate signaling(21, 22, 24, 61). These various treatments were also shown to induce expression of *P2K1*. In order to investigate whether *ILK5* is also strongly induced by these treatments, GUS activity was observed in response to various stresses in transgenic plants expressing the *ILK5promoter::GUS* construct. GUS activity driven by *ILK5* promoter was strongly up-regulated by treatment with eATP, pathogen *Pst* DC3000, the pathogen associated molecular pattern (PAMP) flagellin peptide flg22, MeJA and wounding and relative to mock treatments (*Fig. 3A*, *H* and *J*). These results were confirmed by qRT-PCR analysis using RNA extracted from the treated plants (Fig. 3*D-G*, *I*, *K* and *L*). Similar results were obtained by western blot quantification of *ILK5promoter::GFP* expression using anti-GFP antibody and protein from plants treated with either *Pst* DC3000 or eATP over a time course (*Fig. 3B* and *C*). These results support the notion that both *ILK5* and *P2K1* expression responds to a variety of stresses, consistent with their coordinated role in regulating purinergic signaling initiated by the release to eATP in response to stress.

### ILK5 plays an important role in the pathogen defense response

The biochemical data presented above suggest that defects in ILK5 function should disrupt the normal response to pathogen infection. To examine this directly, we conducted functional analysis using *ilk5* T-DNA insertion mutants (*SI Appendix,* Fig. S7). *ILK5* transcripts were not detected in the *ilk5-1* and *ilk5-3* mutants, suggesting that *ilk5-1* and *ilk5-3* are null alleles. However, in contrast, *ilk5-2* and *ilk5-4* showed low gene expression or no change, likely due to insertion of the T-DNA in the either 5’-UTR or 3’-UTR (*SI Appendix, Fig. S7C* and *D*).

Disease resistance against the virulent pathogen, *luxCDABE*-tagged *Pst* DC3000, was examined with or without addition of ATP. Consistent with our biochemical studies, the *ilk5-1* and *ilk5-3* mutant plants showed enhanced susceptibility to *P. syringae* infection (*Fig. 4A*-*C*). The hyper-susceptible *sid2-2* mutant was used as a positive control(62). These results imply that ILK5 positively regulates pathogen resistance and ATP-inducible defense responses in Arabidopsis. Furthermore, pathogen defense-responsive genes such as *WRKY40* and *CPK28* transcripts were quantified by real-time qPCR under ATP treatment, with both transcripts being significantly reduced in *ilk5-1* and *ilk5-3* mutants (*Fig. 4D* and *E*). According to previous studies, extracellular ATP (eATP) regulates the MAPK pathway through phosphorylation in both mammals and plants(18, 29, 63). To confirm this, we examined the phosphorylation of MPK3/6 in the wild-type and various mutant lines upon treatment with ATP over a time course (*SI Appendix,* Fig. S8). The results showed that *p2k1-1* (D572N; kinase dead), *p2k1-2* (D525N; kinase dead), or *p2k1-3* (T-DNA inserted) mutant plants showed significantly reduced phosphorylation of MPK3/6 compared to wild-type (*SI Appendix, Fig. S8B*). Conversely, phosphorylation of MPK3/6 was significantly increased in transgenic plants ectopically, over-expressing P2K1 (*SI Appendix, Fig. S8C*). P2K1 presumably transmits purinergic signaling via ATP-induced phosphorylation of downstream target proteins to activate the MAPK pathway. In order to understand how ILK5 is involved in MPK3/6 phosphorylation under ATP treatment, phosphorylation of MPK3/6 was examined after ATP treatment. Phosphorylation of MPK3/6 was reduced in both *ilk5-1* and *ilk5-3* mutants (*Fig. 4F*). To confirm these results, phosphorylation of MPK3/6 was restored in complemented lines in which the wild-type *ILK5* gene was driven by the *ILK5* native promoter (*Fig. 4G* and *SI Appendix, Fig. S9A*). Consistent with these findings, we also found that *ilk5-1* plants expressing the ILK5^S192A^ mutant protein were significantly more susceptible to pathogen infection than wild-type or ILK5 complemented plants (Fig. 4*H-J*). In addition, phosphorylation of MPK3/6 was also significantly reduced in *ilk5-1* plants expressing the ILK5^S192A^ protein (*Fig. 4K* and *SI Appendix, Fig. S9B*). These results show that ILK5^S192A^ complemented transgenic plants were more susceptible to pathogen infection and that the ILK5 Ser192 plays an important role in MAPK pathway activation, which impacts the level of disease resistance. To determine if this reduction in phosphorylation of MPK3/6 was not caused by protein destabilization, western blotting was performed using anti-MPK3 or -MPK6 antibodies under ATP treatment in *ilk5* mutants or transgenic plants expressing ILK5^wt^ or ILK5^S192A^ protein in the *ilk5-1* mutant background. Significant MPK3/6 protein reduction or increase was not detected under ATP treatment after 30 minutes, confirming the reduction of phosphorylation was not due to the protein destabilization (*SI Appendix,* Fig. S10). In previous studies, addition of ATP was shown to trigger stomatal closure and enhance innate immunity when plants are sprayed or dip inoculated with the pathogen(8, 64). Hence, it is of interest that *ILK5* is strongly expressed in guard cells and was previously shown to play a role in modulating stomatal aperture(52). Therefore, we examined the ability of the stomata to close in *ilk5-1* mutant plants upon ATP treatment. The movement of the stomata in *ilk5-1* mutant was mis-regulated (*SI Appendix,* Fig. S11). This suggests that, similar to P2K1, ILK5 functions in regulating stomatal closure in response to eATP and, thereby, controls the ability of the bacterial pathogen to enter the leaf via stomata.

**Fig. 4.**
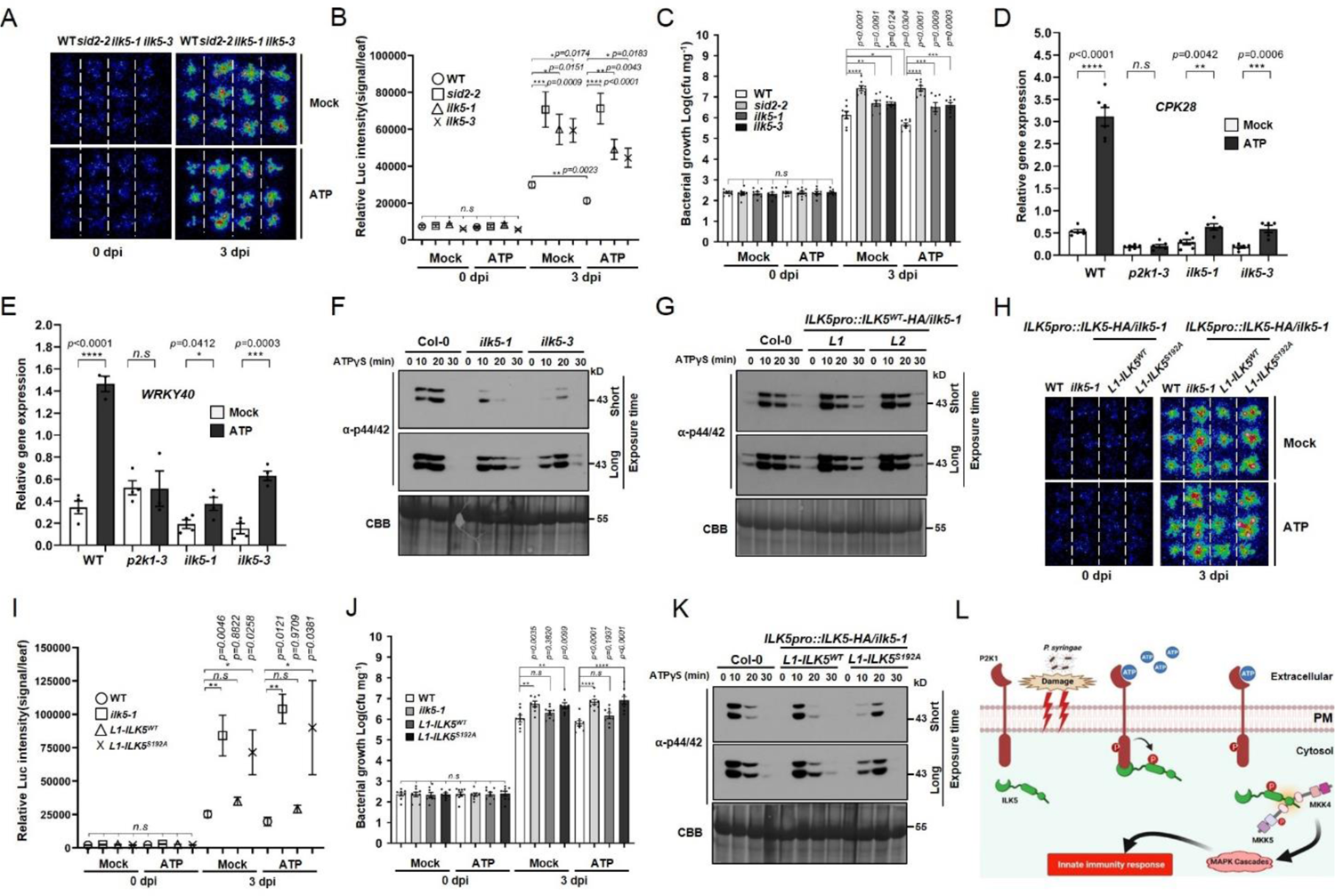
ILK5 is required for plant innate immunity. (*A, H*) Three-week-old plants were flood inoculated with *luxCDABE*-tagged *Pst* DC3000 suspension (OD_600_ = 0.002 approximately 5 × 10^6^ CFU mL^−1^) containing 0.025% (v/v) Silwet L-77 in the presence of ATP (250 μM). Inoculated plants [either *ilk5-1* mutants, complemented *ilk5-1* mutant with ILK5^WT^-HA (L1), or ILK5^S192A^-HA mutant (L1) plants] were used and luciferase luminescence was detected using a low light capture CCD camera either at the time of inoculation (0 days) and 3 days post inoculation. *sid2-2* was used as a control. (*B*, *I*) Quantification of luminescence signal intensities. The signal of *luxCDABE*-tagged *Pst* DC3000 for each plant was monitored, images were captured, and the luciferase signal intensities were quantified using the C-vision/Im32. (*C, J*) Bacterial colonization was determined by plate counting. In panel *B* (n = 9), *C* (n = 8), *I* (n = 6), *J* (n = 8), data shown as mean ± SEM, \*\*\*\**p*<0.0001, \*\*\**p*<0.001, \*\**p*<0.01, \**p*<0.05, *p*-value indicates significance relative to wild-type plants and was determined and analyzed using the GraphPad Prism 8 by one-way ANOVA followed by Dunnett’s multiple comparisons. (*D*, *E*) Real-time qRT-PCR analysis of *WRKY40* and *CPK28* transcripts in *p2k1-3, ilk5-1* and *ilk5-3* mutant backgrounds after ATP (250 μM) treatment. Total RNA was isolated from 2-week-old plants and 2 μg of total RNA was used in this experiment. The *SAND* reference gene was used for data normalization. Data shown as mean ± SD, *WRKY40;* n = 4, *CPK28;* n = 6, \*\*\*\**p*<0.0001, \*\*\**p*<0.001, \*\**p*<0.01, \**p*<0.05, *p*-value was determined and analyzed using the GraphPad Prism 8 by unpaired two-way Student’s *t* test. These experiments were repeated two times (biological replicates) with similar results. (*F*, *G, K*) Phosphorylation of MPK3/MPK6 detected in *ilk5* mutants or complemented lines in which *ILK5* or *ILK5^S192A^* expression was driven by the native promoter by western blotting using anti-phospho 44/42 antibody. ATPγS (250 μM) was added and plants incubated for the times shown. Total protein was extracted at each time point and IB was performed with an anti-44/42 MAPK antibody. CBB staining of protein was used as a loading control. (*L*) Hypothetical model for the role of ILK5 in eATP signaling. Upon addition of the activating ligand eATP, the P2K1 receptor is rapidly auto-phosphorylated and then directly interacts and phosphorylates its downstream target, the ILK5 protein, leading to innate immune response via MAPK cascades. Above the experiments were repeated at least two times (biological replicates) with similar results.

In summary, we hypothesize the following role for ILK5 in plant purinergic signaling. When a pathogen invades causing the release of eATP, the P2K1 receptor is activated by eATP recognition. Subsequently, P2K1 directly interacts with and phosphorylates the downstream ILK5 kinase protein, leading to an innate immune response via activation of the MAPK cascades (*Fig. 4L*).

## Discussion

### Role of Integrin-linked kinase in purinergic signaling

Extracellular ATP is a conserved damaged associated molecular pattern in eukaryotes and, hence, purinergic signaling plays a critical role in the ability of both animals and plants to response to stress(2, 6, 65). Integrins are heterodimeric cell membrane receptors composed of α and β subunits that mechanically function by binding the cell cytoskeleton to the extracellular matrix (ECM)(66). Integrins regulate signal transduction pathways that mediate multiple cellular signals such as organization of cytoskeleton, regulation of the cell cycle, cell adhesion, migration, and activation of growth factor receptors(66–68). Interestingly, integrin and purinergic signaling are closely regulated in metazoans. For example, the human P2Y_2_ nucleotide receptor directly interacts with α_v_β_3_ and α_v_β_5_ integrin complexes via the integrin-binding domain, arginine-glycine-aspartic acid (RGD), contained in P2Y_2_R(69). The RGD sequence was also found to be essential for the role that P2Y_2_R plays in UTP-induced chemotaxis via G_0_ protein coupling. Additionally, Bye et. al., reported that α_IIb_β_3_ (GPIIb/IIIa) integrin protein is activated by P2Y_1_-stimulated Ca^2+^ signaling and P2Y_12_-stimulated activation of the PI3K kinase, which is essential for fibrinogen binding to the platelet of primary rat megakaryocytes(70). Integrins and integrin-linked kinases in animals have been well characterized; however, homologs of animal integrins appear to be absent in plants(71). However, other proteins have been suggested to function like animal integrins in plants. For example, based on the structural similarity to integrin, the Arabidopsis NDR1 protein was suggested to function as an integrin(71). NDR1 was shown to function in plasma membrane-cell wall adhesion and to play an important role in plant innate immunity(71). As the name suggests, integrin-linked kinases strongly interact with the *β* subunits of integrin receptors mediating formation of signaling complexes that are localized at sites of plasma membrane-extracellular matrix interaction (i.e., focal adhesion zones)(46, 55). It is also interesting that P2K1 (LecRK I.9) was reported to mediate plasma membrane-cell wall interaction(21). Hence, it is possible that in the absence of true integrins, plants use such receptors as a means to construct signaling complexes at sites of plasma membrane-cell wall adhesion. Our data would suggest that, similar to metazoans, integrin-like kinases (i.e., ILK5) are likely components of such a signaling complex.

### Phosphorylation of MAPK in purinergic signaling

Activation of a MAPK cascade is a purinergic signaling response shared between animals and plants(72, 73). For instance, P2Y_2_ mediated cell growth is highly stimulated by eATP through MAPK/ERK kinase activation(63, 74). In addition, purinergic signaling activates cPLA2 by another downstream kinase, the mitogen- and stress-activated kinase-1 (MSK1) in order to simultaneously activate both the required ERK1/2 and p38 MAPK(73). Extracellular ATP induces animal cell cycle progression beyond the G1 phase of the cell cycle by activation of P2 receptors(72). When P2 receptors are activated, protein kinase C (PKC) transmits signals to the nucleus through one or more of the MAPK cascades, which possibly include Raf-1, MEK, and ERK, and stimulates transcription factors such as MYC, MAX, FOS, and JUN(6, 72).

Previous research showed that phosphorylation of MPK3/6 was activated in plants upon eATP addition and required P2K1 function primarily but was also reduced in the absence of P2K2(18, 29). However, the mechanism remained unclear as to how eATP recognition at the plasma membrane was coupled to downstream activation of MAPK function. The current study provides this connection by presenting several lines of evidence consistent with the hypothesis that P2K1 activation leads to ILK5 activation, followed by MKK5 activation and then phosphorylation of MPK3/6, the latter as previously published(57). These findings provide new insights into the biological roles and molecular mechanisms of the largely functionally uncharacterized purinergic signaling cascades in plants. Although we found that the Raf-like MAPKKK ILK5 is involved in MPK3/6 phosphorylation, MPK3/6 phosphorylation was not completely abolished in the *ilk5* mutants under ATP treated conditions (*Fig. 4F*) In previous studies, several plant MAPKKKs were reported that regulate the phosphorylation of MPK3/6 in biological processes or environmental stress conditions, especially in innate immunity. For example, Arabidopsis MEKK1 (MEKK1–MKK4/MKK5– MPK3/MPK6) is involved in FLS2 mediated plant immune response(57) and YODA (YDA—MKK4/MKK5– MPK3/MPK6) plays an important role in inflorescence architecture development by promoting localized cell proliferation in Arabidopsis(75) and NPK1 (NPK1–NQK1–NRK1) positively regulates plant cytokinesis during meiosis as well as mitosis in tobacco(76). It was also reported that two highly conserved MAPKKKs, MAPKKK3 and MAPKKK5, mediated MPK3/6 activation in response to activation of several pattern recognition receptors (PRRs) and receptor-like cytoplasmic kinases that play important roles in resistance to several fungal and bacterial diseases in *Arabidopsis*(77). PRRs recognize PAMPs and DAMPs, which then transmit the signals to MAPKKKs, where receptor-like cytoplasmic kinases are heavily involved(78). Many receptor-like cytoplasmic kinases, such as botrytis-induced kinase 1 (BIK1), PBL1, PBL27 and BSK1, are associated with PAMP or DAMP receptors(59, 78, 79). Compared to the MEKK subfamily, Raf-like MAPKKKs are not well characterized in part due to the mutually complementary relationships between the Raf-subfamily of MAPKKK proteins. For instance, the C6 subgroup of RAF-like MAPKKKs, RAF22 and RAF28, are essential for the regulation of embryogenesis in Arabidopsis. More recently, B4 Raf-like MAPKKKs, RAF18, RAF20, and RAF24, were found to interact with and phosphorylate SnRK2s as part of the osmotic stress response(80). There are six ILK members in Arabidopsis, of which ILK4 is the most similar in protein sequence to ILK5. In a recent report, ILK1 was shown to regulate MPK3/6 phosphorylation in response to treatment with the PAMP flagellin flg22 peptide(51). Therefore, ILK5 has the potential to function in a complementary relationship with another subgroup of MAPKKKs or other Raf-like MAPKKKs, including ILK members.

### ILK5 has bona fide kinase activity

Both animal and plant ILKs contain conserved regions consistent with a role as serine/threonine or dual serine/threonine and tyrosine protein kinases. The eukaryotic kinase domain consists of twelve sub-domains with conserved amino acid residues necessary for regulation of kinase activity through binding ATP complexed to divalent cations (e.g., Mg^2+^ or Mn^2+^), the phospho-transfer reaction, and internal residues critical for proper protein folding(81). Animal ILKs contain abundant substitutions in the conserved residues of the kinase domain and, therefore, their functional role as active kinases has been questioned and published reports substantiate the view that they are pseudo-kinases(45). Several members of the plant ILK family, including ILK5, were initially classified as pseudo-kinases because they contain substitutions in some important amino-acid residues thought to be necessary for enzymatic activity(82). However, relative to animal ILKs, plant ILKs contain fewer substitutions and there remained the possibility that they possess kinase function. Indeed, Medicago MsILK1 and Arabidopsis AtILK1 have an unusual Mn^2+^ divalent ion dependent kinase activity, which was found to be necessary for root growth and pathogen response(48, 51). Some of the confusion about ILK kinase function may have to do with the means by which the proteins were purified. For example, in our case with ILK5, and also with the previous report for ILK1, purification of the proteins from plant tissue was necessary to produce proteins with strong kinase activity. In the case of ILK1, its kinase activity is important for mediating plant disease resistance(51).

At present, we have no explanation as to why it is critical to purify these proteins from plant extracts in order to retain kinase activity. It may be that proper protein folding or protein secondary modification in plant tissues may be essential for kinase activation of ILK proteins, including ILK5. In a previous study, according to protein sequence comparison and structural analysis, it was found that ILK4/Raf47, ILK5/Raf27, and ILK6/Raf17 display a relatively longer canonical A-loops and/or C-loops in the sub-domains. Alterations in the canonical kinase motifs can lead to atypical reliance on Mn^2+^ as a cofactor in plant ILKs for full kinase activity(42). Regardless, the data clearly indicate that plant ILKs are functional protein kinases and, hence, appear to function as MAPK kinase kinases to activate downstream MAPK cascades.

### ILK5 plays a key role in regulation of stomatal aperture and plant stress responses

Our previous publication showed that extracellular ATP elicits P2K1-mediated RBOHD phosphorylation to regulate stomatal closure, enhancing plant innate immunity(8). ILK5 is highly expressed in guard cells and strongly induced by various stress conditions and was previously demonstrated to play an important role in regulating stomatal aperture(52)(*SI Appendix*, Fig. 11). It was also shown that mutational loss of MPK3/6 or MKK4/5 function abolished PAMP induced stomatal closure(83). Gain-of-function activation of MPK3/6 induces stomatal closure independently of abscisic acid biosynthesis and signaling(83). This is also true of P2K1-mediated stomatal closure in that *p2k1* mutants still closed stomates in response to ABA treatment(8). The model that emerges from the current study and published results is that purinergic signaling impacts stomatal aperture control, at least in part, via activation of ILK5 and its role in downstream activation of the MAPK cascade.

In addition to playing a role in the plant response to biotic stress, it is also possible that ILK5 is involved in abiotic stress responses. Extracellular ATP accumulates within the apoplast after treatment with such abiotic stress conditions and, interestingly, P2K1 is also highly induced by various abiotic stress conditions (Fig. 3 and *SI Appendix,* S6*E*). The expression of several plant *ILK* genes, including *ILK5,* is increased under various abiotic and/or biotic stress conditions(42), including treatment with bacterial and fungal pathogens, NaCl, heat, cold, and especially wounding, similar to P2K1. Considering that purinergic signaling regulates plant growth under various environmental conditions or in response to diverse abiotic and biotic stress conditions, it seems likely that P2K1 and ILK5 work coordinately to mediate the appropriate plant responses to such stresses. The common feature of these stress responses is their ability to induce the release of eATP.

## Materials and Methods

### Plant materials and growth conditions

*Arabidopsis thaliana* ecotype Columbia (Col-0) and T-DNA insertion mutants, WiscDsLox345-348B17 (*ilk5-1*), SALK_099335C (*ilk5-2*), SALK_050039 (*ilk5-3*) and SALK_082479 (*ilk5-4*) were used in this study. Seeds were sterilized by soaking in 1% bleach solution for 10 min followed by washing five times with sterilized water. Seeds were grown on agar plates for germination. One-half-strength Murashige and Skoog (MS) medium with 1% (w/v) sucrose and 0.5% (w/v) phytagel, 0.05% (w/v) MES, adjusted by KOH to approximately pH 5.7, was used. The plates were then kept in the dark at 4°C for three days for vernalization before being placed in a growth chamber for germination. Seedling plants were transferred to soil and grown to maturity for further experiments. The plants were grown under long-day conditions [16-h days with 150 µE·m^-2^·s^-1^ (E, Einstein; 1 E = 1 mol of photons)]. For harvesting seeds, seedling plants were transferred to soil and grown to maturity at 21°C under long day conditions under 60% relative humidity. Detailed information of plasmid constructions, protein-protein interaction, *in vitro* or *in vivo* phosphorylation assay, RNA extraction and qRT-PCR, MAPK phosphorylation, bacterial growth assay and stomatal closure experiments are provided in *SI Appendix*.

## Data availability

The mass spectrometry proteomics data are available at the ProteomeXchange Consortium via the PRIDE partner repository with the data set identifier PXD006678. All studies are included in the article and/or supporting information.

## Acknowledgments and funding sources

We thank members in Gary Stacey laboratory, especially Samantha Yanders, Sung-Hwan Cho and Jared Ellingsen for aid in writing this manuscript. We thank Scott Peck for providing anti-MPK3 and -MPK6 antibodies. Research was supported by the National Institute of General Medical Sciences of the National Institutes of Health (grant no. R01GM121445), the Next-Generation BioGreen 21 Program Systems and Synthetic Agrobiotech Center, Rural Development Administration, Republic of Korea (grant no. PJ01325403) and through the 3rd call of the ERA-NET for Coordinating Action in Plant Sciences, with funding from the US National Science Foundation (grant 1826803).

## Author Contributions

D.K. designed and performed the experiments; N.A. and J.J.T. performed and supervised the kinase client assay for screening of P2K1 targets. D.Q. performed Co-IP analysis. G.S. supervised the study and edited the manuscript. All authors discussed the results and commented on the manuscript.

## Additional information

This article contains supporting information online at https://

Competing interests: The authors declare no competing interests.

**Supplementary information**

**Supplementary Information Text**

## Extended Description of Materials and Methods

### Plasmid constructs and transgenic plants

Full-length CDS of *ILK5* (AT4G18950), *ILK6* (AT1G14000), *P2K1* (AT5G60300), *RBOHD* (AT5G47910), *CPK5* (AT4G35310), *MKK3* (AT5G40440), *MKK4* (AT1G51660), *MKK5* (AT3G21220), and *MKK8* (AT3G06230), as well as their kinase domain or C-terminal domain, were amplified by PCR reaction using a set of gene-specific primer pairs (Supplementary Table 1). Each PCR product was sub-cloned into *pDONR-Zeo* (Invitrogen) or *pGEM-T* Easy vectors (Promega) for further experiments. In order to generate YFP constructs, *pAM-PAT-GW-YFP* vector was used. For the BiFC experiment in Arabidopsis protoplasts, *pAM-PAT-GW-cYFP*, and *pAM-PAT-GW-cYFP* gateway vector were C-terminally fused with split-YFP. For the LCI assays in *N. benthamiana* leaves, the full-length CDS DNAs from the *pDONR-Zeo* vector were cloned into *pCAMBIA1300-GW-nLUC* and *pCAMBIA1300-GW-cLUC* gateway vector. All gateway clones were generated by LR cloning reactions according to the manufacturer’s instructions (Invitrogen). The *ILK5* promoter was cloned as a 2 kb fragment upstream of the *ILK5* translation start codon into the *pDONR-Zeo* vector by BP reaction. *ILK5 promoter::GUS* and *GFP* fusions were generated by LR cloning into *pGWB3* (GUS) and *pGWB4* (GFP) gateway vectors, respectively. In order to generate the ILK5 wild-type and ILK5^S192A^ complementation lines, *ILK5* genomic DNA (∼4.5 kb), including the 2 kb promoter region, was amplified by PCR reaction using a pair of specific primers and cloned into the *pDONR-Zeo* vector by BP reaction. LR reaction was performed by LR clonase (Invitrogen) with *pGWB13* (HA-tagging) in order to generate the ILK5 wild-type and ILK5^S192A^ complementation lines. Gene expression of *ILK5* was confirmed by qRT-PCR and protein expression by western blotting using HRP conjugated anti-HA antibody (Roche). GST fused-P2K1 and the kinase dead version of P2K1 (GST-P2K1-KD-1; D572N and GST-P2K1-KD-2; D525N) were described by (Chen et. al., 2017)(1) and GST-fused MKK4 and MKK5 were generated by LR reaction into pDEST15 (Invitrogen) gateway vector. In order to generate His-tagged constructs, DNA fragments of *ILK5* and *CPK5* were amplified by PCR reaction and digested with *Eco*RI/*Xho*I and *Bam*HI/*Xho*I, respectively, followed by cloned into the pET21a vector. ILK5^S192A^, ILK5^S192D^, MKK4^K108R^, MKK5^K99R^ and MKK5^K99RT215AS221A^ were generated by site-directed mutagenesis following the manufacturer’s protocol (Invitrogen, Platinum SuperFi DNA polymerase). To obtain transgenic plants, each construct was introduced into *Agrobacterium tumefaciens* strain GV3101 and Arabidopsis plants were transformed using the ‘floral-dip’ method(2). The transgenic plants were selected by germination on 20 mg/L hygromycin and 100 mg/L cefotaxime-containing one-half-strength MS medium with 1% (w/v) sucrose and 0.5% (w/v) phytagel and 0.05% (w/v) MES adjusted by KOH to approximately pH 5.7 (Duchefa, Haarlem, The Netherlands) under long-day conditions at 21°C. Homozygous plants were obtained by self-pollination (T3 generations) and confirmed by hygromycin selection.

## Kinase client assay (KiC assay)

The KiC assay was performed as previously described(8, 54). Briefly, a library of more than 2100 peptides developed from identified *in vivo* phosphorylation sites taken from a number of studies was incubated with the purified recombinant GST fused P2K1 kinase domain (GST-P2K1-KD) in the presence of ATP. Two sets of empty vectors (GST and MBP) and two kinase-dead proteins, GST-P2K1-KD-1 (D572N) and GST-P2K1-KD-2 (D525N), were used as negative controls. The peptide reaction mixtures were analyzed using a Finnigan Surveyor Liquid Chromatography system interfaced with a LTQ Orbitrap XL ETD mass spectrometer. For final validation, each spectrum was manually examined and the phosphopeptides were allowed only if the highest pRS site probability, pRS score, Xcorr value, and site-determining fragment ions were observed for the unambiguous location of the phosphorylation site. Phosphopeptides with a pRS score of ≥15 and/or a pRS site probability of ≥50% were accepted.

## Immunoblot assay

Total protein was extracted from 10-day-old Arabidopsis transgenic plants at the indicated time points by homogenization in extraction buffer containing 50 mM Tris-HCl (pH 7.5), 150 mM NaCl, 10 mM MgCl_2_, 1 mM EDTA, 1 mM DTT, 0.2 mM Phenylmethylsulfonyl fluoride (PMSF) (Sigma, 93482), 10% glycerol, 0.5% Triton-X 100, and 1 x protease inhibitor (Sigma Aldrich, PIA32955) at 4°C with gentle agitation for 2 h to extract total protein from the plant tissues. The samples were centrifuged at 20,000 x g for 15 min at 4°C. Supernatant was transferred to new tube and centrifuged again at 20,000 x g for 10 min at 4°C, to pellet any carryover leaf debris. The extracted total proteins were mixed with 5 x Laemmli loading buffer containing 10% SDS, 50% glycerol, 0.01% bromophenol blue, 10% *β*-mercaptoethanol, 0.3 M Tris-HCl pH 6.8, and heated in boiling water for 5 min. The total extracted proteins were separated by 12% SDS-PAGE gel and detected by immunoblotting with anti-HA-HRP (Roche, 12013819001; dilution 1:2000), anti-GFP (Invitrogen, A11122; dilution 1:4000), anti-MPK3 (dilution 1:2000) and MPK6 (dilution 1:4000) antibody(84, 85).

## Protoplast isolation and DNA transformation

Full-length *ILK5* and *P2K1* CDS for the YFP-fused constructs were prepared by PCR reaction and fused to the 5′-end of YFP in the *pAM-PAT-GW-YFP* vector for transient expression and stable transformation. For the transient expression of YFP fused to ILK5 or P2K1, the YFP-fused constructs were introduced by polyethylene glycol (PEG)-mediated transformation into Arabidopsis protoplasts prepared from leaf tissues of three- to four-week-old plants. Before the isolation of Arabidopsis protoplasts, the plants were incubated with 1 M mannitol for 30 min. 30 mL of 0.22 μm filter-sterilized enzyme solution was added containing 10 mM MES-KOH (pH 5.7), 0.4 M mannitol, 1 mM CaCl_2_, 1% cellulase (Onozuka R-10), 0.25% macerozyme (R-10) and 1% BSA (Goldbio) and 0.035% *β*-mercaptoethanol, covered with parafilm and aluminum foil for dark conditions. After gentle agitation at 21°C for 8-10 h, the enzyme solution containing protoplasts was gently filtered through the 75 mm nylon mesh into a 50 mL tube. These protoplasts were gently covered with 10 mL W5 solution (2 mM MES pH 5.7, 154 mM NaCl, 125 mM CaCl_2_, and 5 mM KCl) without disturbing the 21% sugar content gradient, followed by centrifugation for 8 min at 100 x g. Approximately 10 mL of intact protoplasts floating on the sucrose to 20 mL of W5 solution were carefully transferred to a new 50 mL tube. An aliquot of 15 mL W5 solution was added followed by centrifugation for 5 min at 60 x g. The protoplasts were washed with 15 mL of W5 solution and centrifuged again for 5 min at 60 x g. Pelleted protoplasts were resuspended in 5 mL MaMg Solution containing 4 mM MES pH 5.7, 0.4 M mannitol and 15 mM MgCl_2_. Plasmids encoding the proteins to be expressed were transfected into protoplasts using the plasmid-PEG-calcium transfection method(3, 4). Briefly, 20 μg of each construct (1 μg/μL) was added to 300 μL protoplast solution and mixed gently. 320 μL of PEG solution were added and gently mixed with DNA-protoplasts by tapping. This mixture was incubated at 21°C for 30 min. The transfection mixture was diluted and washed with 1 mL W5 solution 5-times at 5 min intervals, then centrifugated at 50 × g for 4 min at room temperature. The protoplasts were resuspended gently with 2 mL W5 solution. After incubation for 24 h under dark conditions, 200 μM ATP was added to the protoplast solution for further experiments.

## Subcellular localization of YFP-tagged protein and bimolecular fluorescence complementation (BiFC) assay

YFP or Split-YFP protein fusion plasmids (*pAM-PAT-GW-YFP* or *pAM-PAT-YFP-GW* for YFP and *pAM-PAT-GW-nYFP* and *pAM-PAT-GW-cYFP* for split-YFP) were co-transformed into Arabidopsis protoplasts as described above and then incubated in a 21°C growth chamber for 24 h under dark conditions. The YFP fluorescence was monitored using a Leica DM 5500B compound microscope with Leica DFC290 color digital camera 24 h after transformation. 5 μM FM4-64 (Invitrogen, T3166) was used as a counter stain for the plasma membrane. In addition, 1 μg/mL of 4′-6-diamidino-2-phenylindole (DAPI; Sigma-D9542-10MG) was used as a nuclear marker. For the YFP transgenic plants, YFP constructs (driven by the *35S* promoter) were transformed into the *A. tumefaciens* strain GV3101 (pMP90RK) and Arabidopsis plants were transformed using the ‘floral-dip’ method(2). The transgenic plants were selected by germination on 20 μg/mL hygromycin-containing MS medium under long-day conditions at 21°C. Homozygous T3 generation plants were selected after self-pollination. Confocal images were generated using a laser confocal microscope (Leica, TCS SP8 STED) attached to a vertical microscope (Leica MP Color Digital Camera) equipped with various fluorescein filters. The YFP signal was excited at a wavelength of 514 nm under a confocal laser-scanning microscope with an argon ion laser system. The fluorescence images were collected in the YFP or FM4-64 channels. 30 μL monoclonal anti-HA Agarose beads (Sigma, A2095-1 mL), spun down and washed seven times with extraction buffer.

## Co-immunoprecipitation assay

*A. tumefaciens* GV3101 carrying the indicated constructs in infiltration buffer [10 mM MES (pH 5.7), 10 mM MgCl_2_, 100 μM 4’-Hydroxy-3’,5’-dimethoxyacetophenone] was infiltrated into 4-week old leaves of *N. benthamiana*. Total protein was purified from co-infiltrated *N. benthamiana* leaf tissues using the protein extraction buffer: 50 mM Tris-HCl (pH 7.5), 150 mM NaCl, 0.2 mM PMSF, 0.5% Triton-X 100, and 1 x protease inhibitor (Thermo Fisher Scientific; A32955) by gentle agitation at 4°C for 1 hours. The solution was centrifuged at 20,000 x g for 15 min at 4°C. The supernatant was decanted in a new e-tube and 30 μL monoclonal anti-HA antibody Agarose beads (Sigma, A2095-1 mL) was added, and incubated overnight with end-to-end shaking at 4°C. Subsequently, washed at least 7 times with washing buffer containing 50 mM Tris-HCl (pH 7.5), 150 mM NaCl, and 1 x protease inhibitor. After washing, the resin was eluted with 25 μL 1 x SDS-PAGE loading buffer and the eluent heated in boiling water for 10 min. The proteins were separated by 10% SDS-PAGE gel electrophoresis and detected by immunoblotting with anti-HA-HRP (Cat no. 12013819001, Roche; dilution, 1:1000) a horseradish peroxidase (HRP)-conjugated anti-HA and anti-Myc (Cat no. SAB4700447, Sigma-Aldrich; dilution, 1:2000) antibodies. MKK8-myc was used as a negative control.

## Split-luciferase complementation imaging (LCI) assay

Each of the constructs containing full-length cDNAs were cloned with N- or C-terminal LUC (nLUC or cLUC) at their C-termini of the *pCAMBIA1300-GW-nLUC* or *pCAMBIA1300-GW-cLUC* vector by LR cloning and transformed into the *A. tumefaciens* GV3101 by GenePluser^TM^ electro-transformation (BioRad). The positive strains were selected by Rifampicin (25 μg/mL) and Kanamycin (50 μg/mL) antibiotics and then cultured in Luria-Bertani (LB) medium. When the OD_600_ value of the cultures reached 1.0, the cultures were resuspended with infiltration buffer [10 mM MES (pH 5.7), 10 mM MgCl_2_, 100 μM 4’-Hydroxy-3’,5’-dimethoxyacetophenone] and incubated for 2 h. *A. tumefaciens* GV3101 carrying the indicated constructs (OD_600_ = 0.6) was infiltrated into 4-week old leaves of *N. benthamiana*. The infiltrated leaves were incubated at 28°C for 2 days before LUC activity measurement. 1 mM D-luciferin containing 0.01% silwet-L77 was sprayed onto the *N. benthamiana* leaves and immediately placed in dark conditions for 10 min to quench the fluorescence. The luminescence was monitored and captured using a low light imaging CCD camera (Photek; Photek, Ltd.).

## GST-, His recombinant protein expression and purification

His-tagged ILK5 and CPK5 were fused C-terminally in the *pET21a* vector (Novagen). The GST-tagged intracellular domain of P2K1 (GST-P2K1-KD) and, its kinase dead versions (GST-P2K1-KD-1; D572N) and GST-P2K1-KD-2; D525N) were fused N-terminally in the *pGEX-5X-1* vector (GE Healthcare). Plasmids were transformed into Rosetta^TM^ (DE3) competent cells (Novagen) expressing YopH tyrosine phosphatase. Bacterial cultures in LB medium were induced with 0.5 mM isopropyl *β*-D-1-thiogalactopyranoside (IPTG) after reaching an OD_600_ absorbance of 0.5 and incubated at 28°C for an additional 4 h. Bacterial cells were collected at 6,000 x g for 10 min and His- and GST-tagged proteins were purified by TALON Metal Affinity Resin (Clontech #635502) and Glutathione Resin (GenScript #L00206) following the manufacturer’s protocol, respectively. For the His-tagged ILK5 protein from *N. benthamiana*, CDS of *ILK5* was amplified by PCR and digested with *Sma*I/*Xho*I restriction enzyme and ligated into the *pEAQ-HT* vector(5). These constructs were introduced into *A. tumefaciens* strain LBA4404 and infiltrated into leaves from 4-6 weeks old *N. benthamiana*. In order to extract the His-tagged ILK5 protein, we followed the previous protocols(5, 6).Leaves were harvested 3-5 days after infiltration of *N. benthamiana* and flash-frozen in liquid nitrogen. Harvested leaves were ground in a mortar and pestle with liquid nitrogen until a superfine powder. 50 mL total protein extraction buffer [50 mM sodium phosphate (pH 8), 10 mM imidazole, 0.5 M NaCl, 1% Triton-X 100, and 1 x protease inhibitor] were added in each sample, then incubated at 4°C with gentle agitation for 2 h to extract total protein from the infiltrated *N. benthamiana*. Plant debris was pelleted by centrifugation at 4°C and 20,000 x g for 20 min. Supernatant was transferred to a new tube and centrifuged again at 4°C and 20,000 x g for 10 min to pellet any carryover leaf debris. The supernatant was transferred to new tubes repeatedly until no debris could be detected. His-tagged ILK5 protein was purified by TALON Metal Affinity Resin (Clontech #635502) following the manufacturer’s protocol. His-tagged ILK5 protein was eluted by Elution buffer [50 mM sodium phosphate (pH 8.0), 300 mM NaCl, 150 mM imidazole and 1 x protease inhibitor]. All purified protein was concentrated up to approximately 1 μg/μL and contaminants removed for further experiments using Vivaspin 2 or 6 (30 kDa MWCO, GE Healthcare) columns following the manufacturer’s protocol. The protein concentration was measured with the BIO-RAD Quick Start Bradford Dye Reagent and confirmed by the NanoDropOne Spectrophotometer.

## *In vitro* kinase assays

*in vitro* kinase assay was performed as previously described with minor modifications(7). Briefly, 2 μg of purified GST or GST-tagged protein kinases were incubated with 2 μg His-tagged ILK5 protein as a substrate in a 25 μL reaction buffer containing 20 mM Tris-HCl (pH 7.4), 5 mM MgCl_2_ or MnCl_2_, 100 mM NaCl, 2 mM ATP, with or without 0.2 μL radioactive [γ-^32^P] ATP for 3 h at 30°C. The two reactions with or without radioactive [γ-^32^P] ATP reactions were stopped by adding 5 μL of 5× SDS loading buffer and incubating in the thermomixer (Eppendorf 22331 Hamburg) at 100°C for 5 min. Each reaction was separated by electrophoresis in 12% SDS-PAGE gels, and the gel containing radioactive [γ-^32^P] ATP was auto-radiographed using a Typhoon FLA 9000 Phospho-imager (GE Healthcare) for 24 h. Either reaction mixture with or without radioactive [γ-^32^P] ATP were stained with CBB (Coomassie Brilliant Blue) and used as a loading control. MBP (Myelin Basic Protein) was used as a generic substrate and CPK5 and GST were used as negative controls, respectively.

## *β*-glucuronidase (GUS) assay

Whole seedlings or various tissues were immersed in histochemical staining solution containing 1 mM 5-bromo-4-chloro 3-indolyl *β*-glucuronic acid, 100 mM sodium phosphate (pH 7.0), 0.1 mM EDTA, 0.5 mM ferricyanide, 0.5 mM ferrocyanide and 0.1% Triton-X 100. After incubation in a vacuum for 10 min, the seedlings were incubated at 37°C for 6–12 h depending on staining status. For the wounding treatment, rosette leaves were wounded using hemostat forceps. Chlorophyll was cleared from the plant tissues by immersing them in 70% ethanol then washing with 70% EtOH repeatedly until tissue was clear. Stained tissues were observed by Fisher Stereo-master microscope (FW02-18B-1750) and digital images were obtained using an AmScope digital camera (MU1000).

## RNA extraction and qRT-PCR analysis

Two-week-old seedling plants were incubated in sterile one-half-strength liquid MS at room temperature overnight. Samples were collected after treatment with 200 μM ATP, 100 nM flg22, *luxCDABE*-tagged *P. syringae DC3000*, drought, heat (37°C), cold (4°C), NaCl (200 mM) or wounding. Plants were immediately harvested by flash-freezing in liquid nitrogen. Harvested leaves were ground in a mortar and pestle with liquid nitrogen until a superfine powder. Total RNA was extracted using Trizol reagent (Invitrogen, 15596018) according to the manufacturer’s instructions. The RNA concentration was estimated followed by treatment with Turbo DNA-free DNase (Invitrogen, AM2238). The RNA was used for first-strand cDNA synthesis using reverse transcriptase (Promega, M-MLV Reverse Transcriptase, PRM1705). The real-time PCR was performed using the SYBR^TM^ Green PCR Master mix (Applied Biosystems, Power SYBR® Green PCR Master Mix, 4368702) following the manufacturer’s instructions. The gene-specific primers used are listed in the *SI Appendix* Table S1. RNA levels were normalized against the expression of the *SAND* or *UBIQUITIN* (*UBQ*) gene.

## *In vivo* kinase assay

100 μg of pUGW-CPK5-HA or pUGW-ILK5-HA(WT or S192A) constructs were introduced by polyethylene glycol (PEG)-mediated transformation into Arabidopsis protoplasts prepared from leaf tissues of three- to four-week-old plants described above, then incubated in a 21°C growth chamber for 48 h under dark conditions with 2 mL W5 solution (2 mM MES pH 5.7, 154 mM NaCl, 125 mM CaCl_2_, and 5 mM KCl). 250 μM ATP or ATPγS (poorly hydrolyzed ATP analog) was added and incubated for 1 hour at 21°C. All intact protoplasts were then subsequently harvested by centrifugation at 50 × g for 4 min at room temperature. All supernatant was removed and 250 μL of total protein extraction buffer [50 mM Tris-HCl (pH 7.5), 150 mM NaCl, 0.2 mM PMSF, 0.5% Triton-X 100, and 1 x protease inhibitor (Thermo Fisher Scientific; A32955)] was added and incubated 4°C for 1 hour. Immunoprecipitations were performed by 30 μL monoclonal anti-HA Agarose beads (Sigma, A2095-1 mL), spun down and washed seven times with extraction buffer. After washing, 50 μL 1x SDS-PAGE loading buffer was added and heated at 100°C for 10 min. The proteins were separated by SDS-PAGE and western blotting was carried out using Anti-HA-HRP (Roche, 12013819001; dilution 1:2000) and anti-phospho-Ser/Thr (BD Transduction laboratories, 612548, dilution 1:6000) antibodies using the method described above.

## MAPK phosphorylation assay

Leaf discs from 4- or 5-week-old plants were incubated in ddH_2_O overnight at 21°C, then treated with 200 μM ATP or ATPγS, for 0, 10, 20, or 30 min. Leaf discs treated with ATP were harvested at each time point by flash-freezing in liquid nitrogen. Harvested leaves were ground in a mortar and pestle with liquid nitrogen. Total protein was extracted with extraction buffer containing 50 mM Tris (PH 7.5), 150 mM NaCl, 0.5% Triton-X 100, and 1 x protease inhibitor at 4°C with gentle agitation for 2 h to extract total protein from the leaf discs. The samples were centrifuged at 20,000 x g for 15 min at 4°C. Supernatant was transferred to new tube and centrifuged again at 20,000 x g for 10 min at 4°C, to pellet any carryover leaf debris. The extracted total proteins were separated by 10% (w/v) SDS-PAGE gel and detected by immunoblotting with anti-phospho-p44/p42 MAPK antibody (Cell signaling, 9102S; dilution, 1:2000).

## Bacterial growth assays

Two or three week-old plants were flood inoculated with a suspension of *luxCDABE*-tagged *P. syringae DC3000* cells (OD_600_ = 0.002 approximately 5 × 10^6^ CFU mL^−1^) containing 10 mM MES pH 5.7, 10 mM MgCl_2_ and 0.025% (v/v) Silwet L-77 with or without the addition of 250 μM ATP. The bacterial suspension was immediately removed by decantation and the plates containing inoculated plants were incubated in the plant chamber under long-day conditions (16-h days with 150 µE·m^-2^·s^-1^) at 21°C. Three days post inoculation, plants were washed with ddH_2_O for 5 min. The plants without roots were ground in 10 mM MgCl_2_, diluted serially, and plated on LB agar with 25 mg/L rifampicin. Colonies (CFU) were counted after incubation at 28°C for 2 days.

## Stomatal closure experiment

Four-week-old plants were placed under light for 2 h in order to ensure that the starting plants had fully open stomata. Leaf peels were obtained from the abaxial side by tweezers and incubated in a stomata buffer containing 10 mM MES (pH 6.15), 10 mM KCl, and 10 μM CaCl_2_. These peels were treated with 2 mM ATP, 5 μM ABA (Sigma, A1049) or mock solution for 1 h and then images captured using a light microscope (Nikon, Alphaphot2). The stomatal aperture was measured using ImageJ software (Version 1.51a).

---

**Fig. S1.**
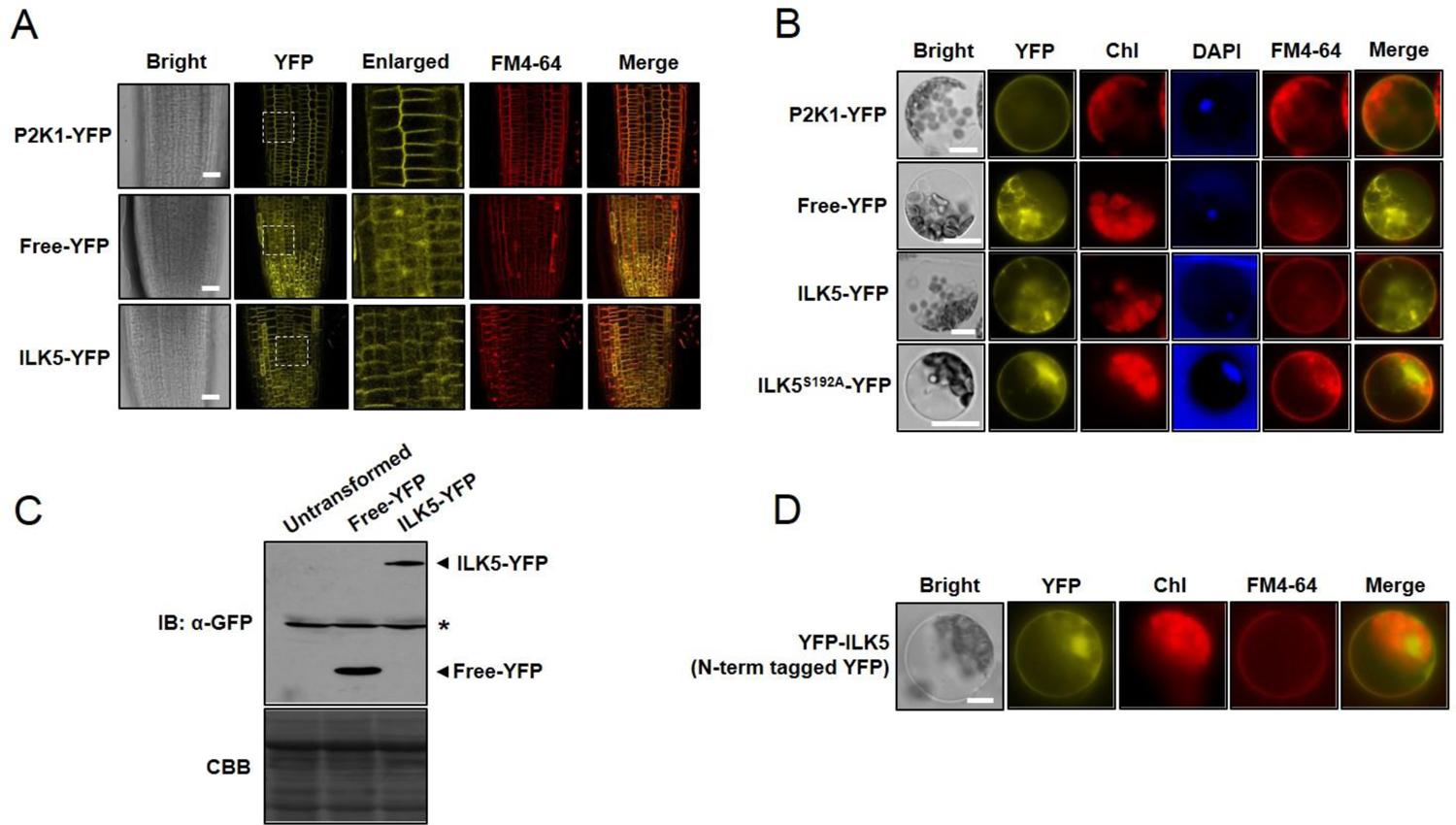
Subcellular localization of ILK5 protein. (*A*) Subcellular localization of ILK5 in plants expressing a *35S::ILK5*-*YFP* construct. Fluorescent confocal images displaying the subcellular distribution of ILK5-YFP protein were detected from primary root tissue of seven-day-old seedlings. Plasma membranes were counter-stained by incubation for 1 min in a FM4-64 solution (5 μM). Images were collected by spinning-disk fluorescence confocal microscopy using a Leica 40 x Plan-Apochromat lens with appropriate EYFP and PI fluorescence filter sets. Dotted square line indicates enlarged area of the YFP images. P2K1-YFP and free-YFP were used as controls. FM4-64 was used as a plasma membrane marker. Merge indicates overlapped image YFP and FM4-64. Scale bars = 50 μm. (*B*) Subcellular localization of ILK5 and ILK5^S192A^ proteins in Arabidopsis protoplasts. YFP-fused constructs were introduced by polyethylene glycol (PEG)-mediated transformation into Arabidopsis protoplasts prepared from three-week-old wild-type Arabidopsis plants and incubated for 24 h under dark conditions. Scale bars, 10 μm. The YFP fluorescence was monitored using a Leica DM 5500B Compound Microscope with Leica DFC290 Color Digital Camera 24 h after transformation. DAPI and FM4-64 were used as nuclear and plasma membrane markers, respectively. Chl represents chlorophyll auto-fluorescence signal. Merge indicates overlapped image of YFP and FM4-64. P2K1-YFP localizes to the plasma membrane and Free-YFP is diffused within the cytosol and nucleus, respectively. Scale bars: 10 μm. (*C*) Western blotting analysis of ILK5-YFP. ILK5-YFP protein was extracted from tissues (as shown in panel B) and used for western blotting using anti-GFP antibody. Total protein was extracted from untransformed tissues and those expressing either free-YFP or ILK5-YFP driven by the *35S* promoter. CBB staining of protein was used as a loading control. Asterisk represents non-specific bands. Note that no proteolysis of the ILK5-YFP protein was detected. (*D*) Subcellular localization of YFP-ILK5 (N-terminally tagged YFP). FM4-64 was used as a plasma membrane marker. All above experiments were repeated three times with similar results.

**Fig. S2.**
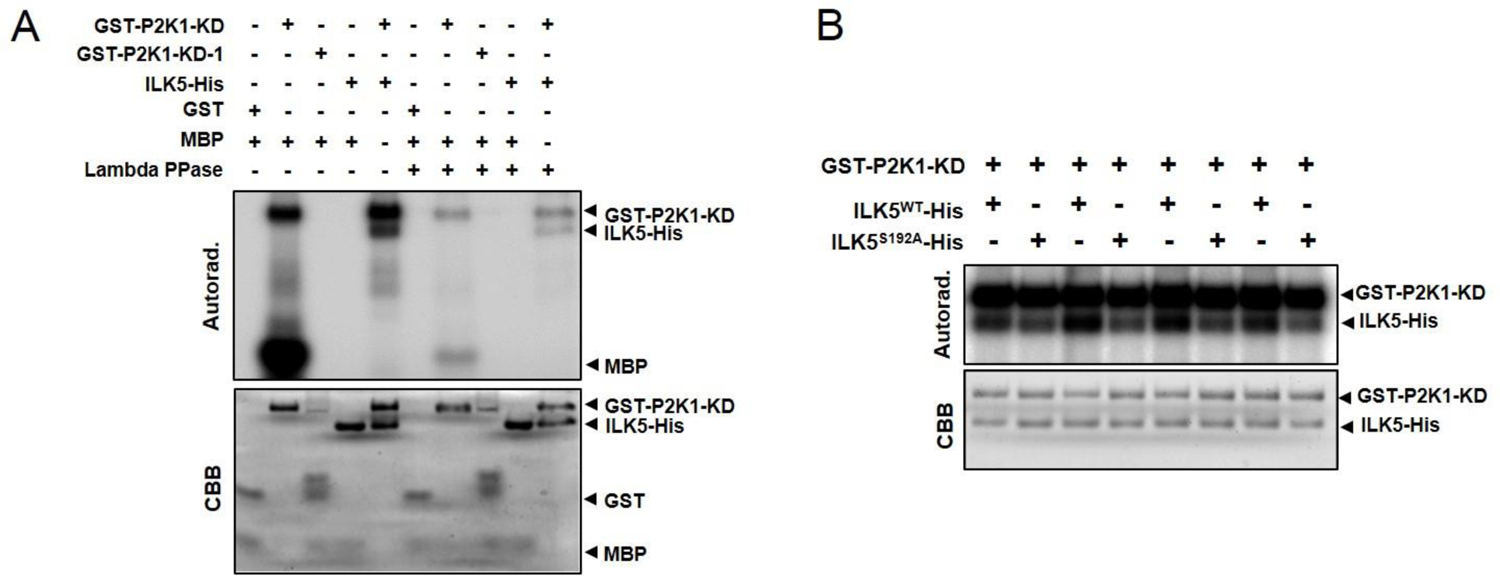
P2K1 directly phosphorylates ILK5. (*A*) Confirmation of ILK5 phosphorylation by treatment with Lambda protein phosphatase (Lambda PPase). Lambda PPase was added to release phosphate groups from phosphorylated serine, threonine, and tyrosine residues. Auto- and trans-phosphorylation were detected by incorporation of γ-[^32^P]-ATP. MBP and GST were used as a universal substrate and a negative control, respectively. Protein loading was visualized by coomassie brilliant blue (CBB) staining. This experiment was repeated three times with similar results. (*B*) P2K1 phosphorylates ILK5 at Ser192 residue. Purified recombinant ILK5-His and ILK5^S192A^ proteins were incubated with GST-P2K1 kinase domain (GST-P2K1-KD) for *in vitro* kinase assay. Autophosphorylation and trans-phosphorylation were detected by incorporation of γ-[^32^P]-ATP. The protein loading was visualized by CBB staining. The intensity of each band was measured by autoradiograph using Image j software (1.52a) and designated in *Fig. 1J*.

**Fig. S3.**
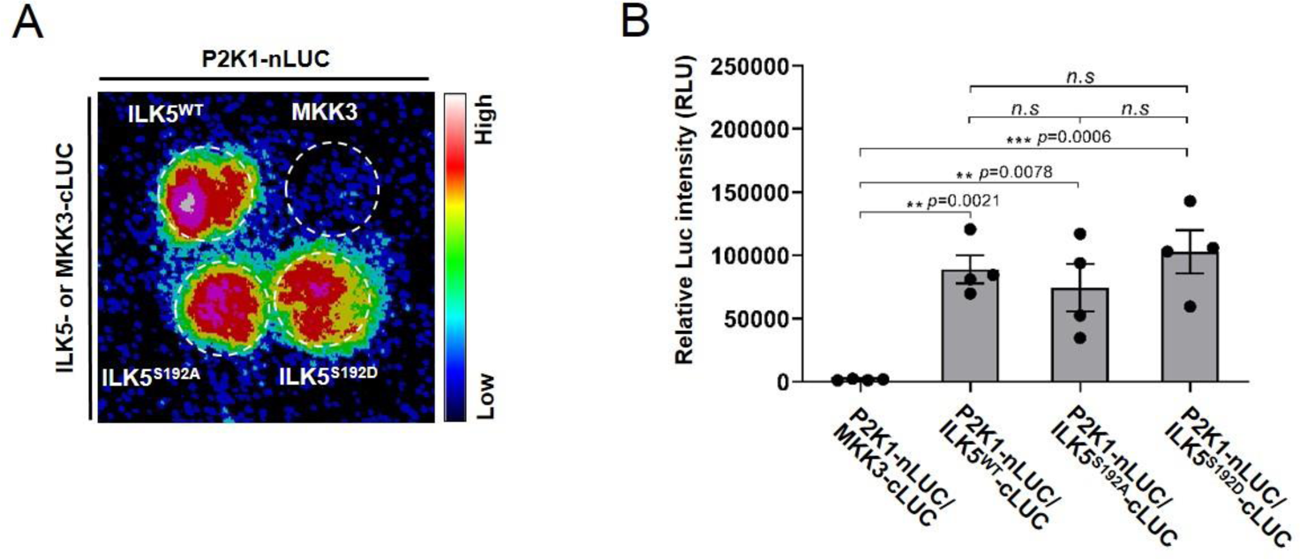
Demonstration of P2K1-ILK5^S192A^ and P2K1-ILK5^S192D^ mutant protein interactions. (*A*) Demonstration of P2K1-ILK5^S192A^ and P2K1-ILK5^S192D^ mutant protein interactions by split-luciferase imaging assay. MKK3 was used as a negative control. (*B*) Quantification of P2K1-ILK5 mutation versions interaction signal intensities. P2K1-ILK5 interaction was monitored, images were captured, and the luciferase signal intensities were quantified using the C-vision/Im32 and analyzed using the GraphPad Prism 8 program. Data shown as mean ± SEM, n = 4 (biological replicates), \*\*\*\**p*<0.0001, *** *p*<0.001, ** *p*<0.01, * *p*<0.05, *p*-value indicates significance relative to MKK3-cLUC and was determined by one-way ANOVA followed by Dunnett’s multiple comparisons. All above experiments were repeated three times with similar results.

**Fig. S4.**
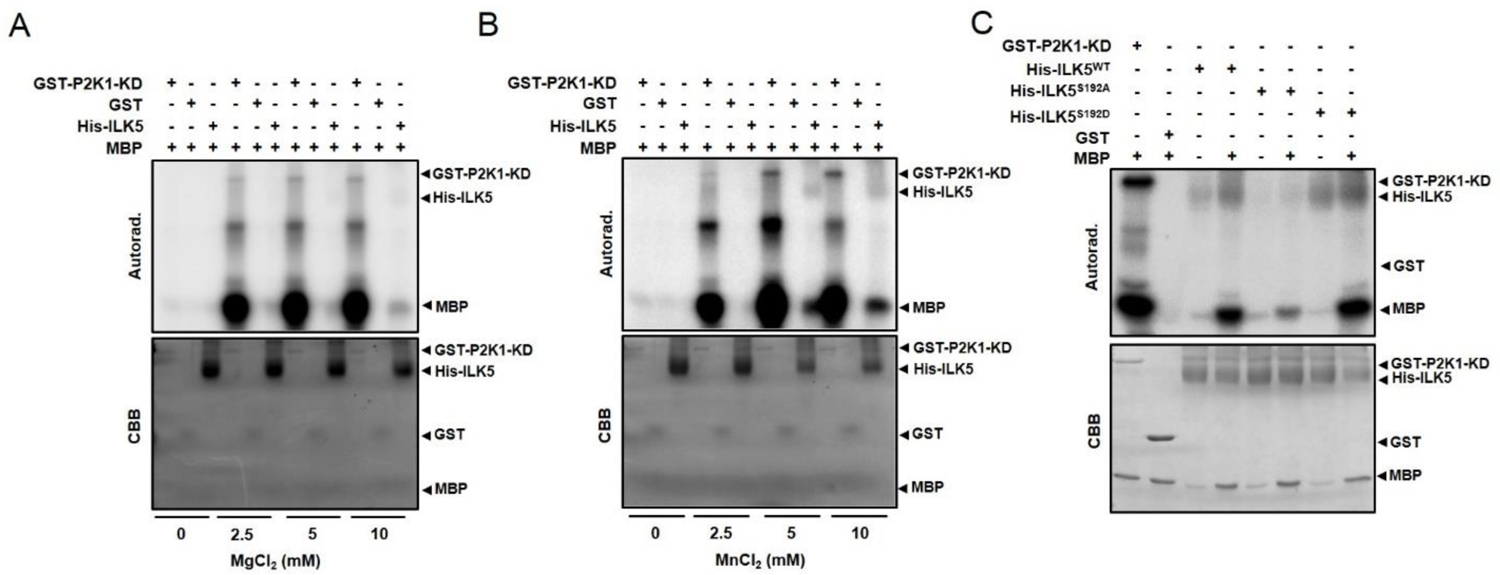
(*A, B*) ILK5 has bona fide kinase activity. Purified recombinant His-ILK5 protein extracted from *N. benthamiana* leaves was incubated with MBP for *in vitro* kinase assay. Autophosphorylation and trans-phosphorylation were detected by incorporation of γ-[^32^P]-ATP with the indicated concentrations of MgCl_2_ or MnCl_2_. (*C*) *in vitro* kinase assay of purified recombinant His-ILK5 protein (WT, S192A and S192D) extracted from *N. benthamiana*. P2K1 and GST were used as a positive and negative controls, respectively. MBP was used as a universal substrate. The protein loading was visualized by CBB staining. All above experiments were repeated two times with similar results.

**Fig. S5.**
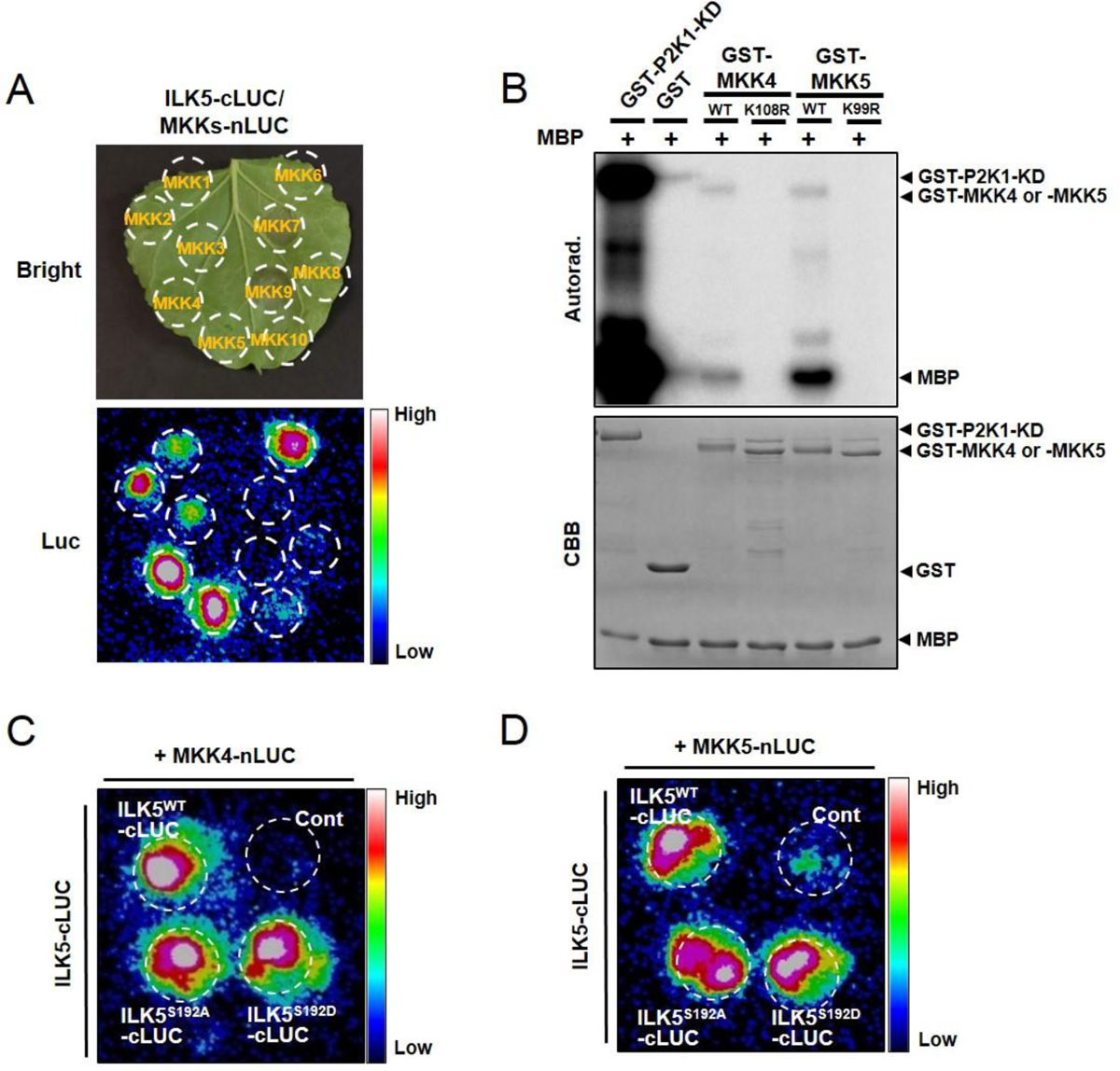
LCI analysis of ILK5 putative interactors among MKKs and investigation of MKK4, MKK5 kinase activity. (A) ILK5 interaction between MKKs (1–10) was demonstrated by split-luciferase imaging. Split-luciferase imaging was performed in *N. benthamiana* leaves co-infiltrated with *A. tumefaciens GV3101* strain expressing ILK5 and MKKs. Dotted circles indicate the infiltrated area in *N. benthamiana* leaves. Note the very strong interaction with MKK4 and MKK5. (*B*) Purified wild-type recombinant proteins, GST-P2K1-KD, GST, GST-MKK4, GST-MKK5 or kinase dead versions, GST-MKK4^K108R^ and GST-MKK5^K99R^, were incubated with MBP for *in vitro* kinase assay. Autophosphorylation and trans-phosphorylation were detected by incorporation of γ-[^32^P]-ATP. P2K1 and GST were used as positive and negative controls, respectively. MBP was used as a universal substrate. The protein loading was visualized by CBB staining. (*C* and *D*) Demonstration of ILK5^S192A^ or ILK5^S192D^ and MKK4 or MKK5 interaction by split-luciferase imaging assay. Cont represents co-infiltrated region of ILK5-cLUC with MKK8-nLUC. It was used as a negative control. Above experiments were repeated three times with similar results.

**Fig. S6.**
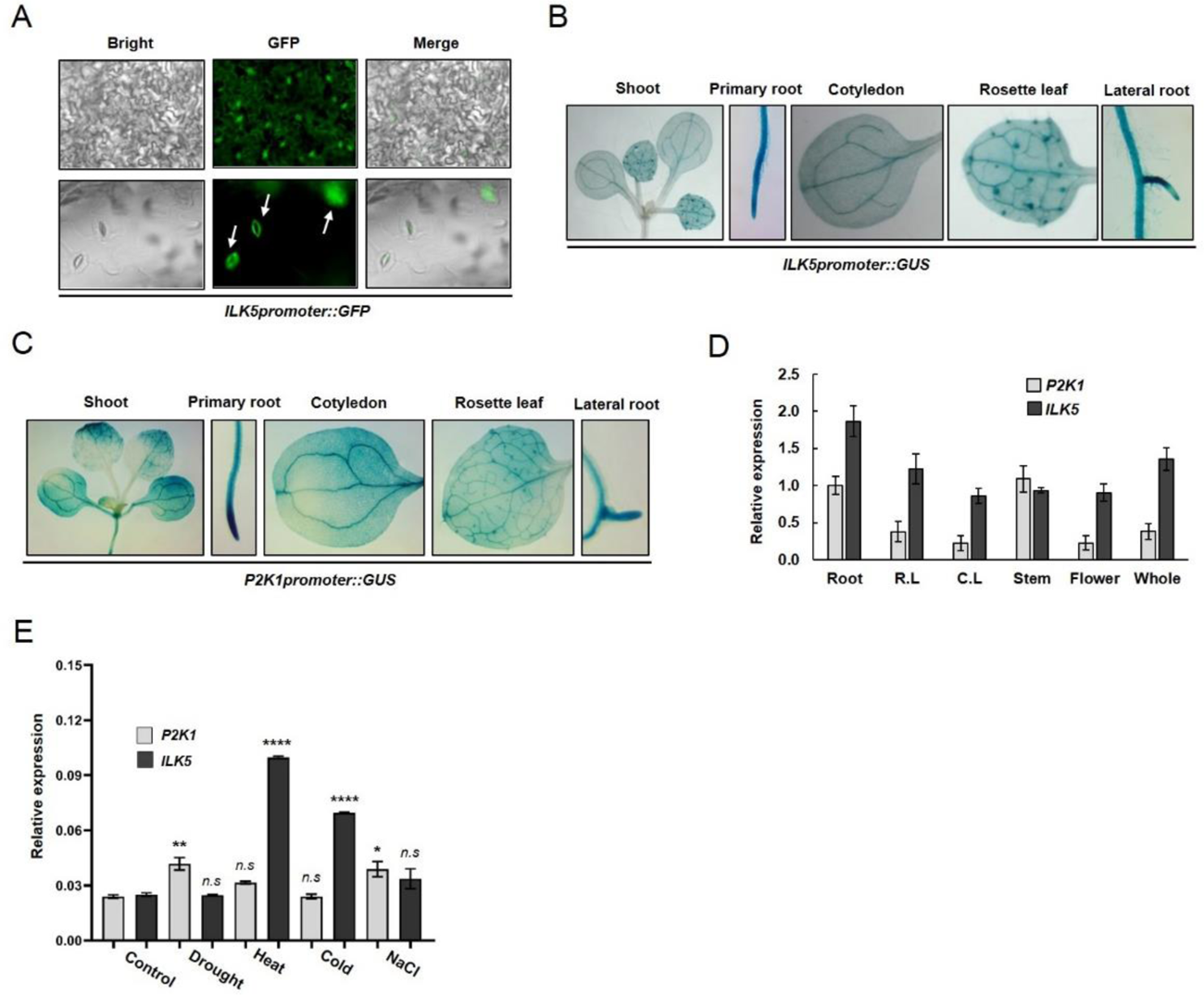
*ILK5* and *P2K1* are ubiquitously expressed in various plant tissues. (A) *ILK5* is highly expressed in guard cells. *ILK5promoter::GFP* transgenic plant was constructed in wild-type Arabidopsis plants transformed with a chimeric *ILK5promoter::GFP* [i.e., 2 kb of the *ILK5* promoter (5′ region of the *ILK5* gene) fused to the GFP coding sequence]. Fluorescent confocal images displaying the GFP expression were taken from rosette leaf tissue of 10 day-old seedlings. The YFP fluorescence was monitored using a Leica DM 5500B Compound Microscope with Leica DFC290 Color Digital Camera. Arrows indicate guard cells expressing GFP. Merge indicates overlapped image Bright and GFP. (*B* and *C*) Histochemical analysis of *ILK5pro::GUS* and *P2K1pro::GUS* expression in various tissues. The expression patterns in plants expressing either *ILK5promoter::GUS* or *P2K1promoter::GUS* transgenic plant. transgenic plant were detected by histochemical staining in 10 day-old seedling grown under normal conditions. (*D*) Real-time quantitative RT-PCR (qRT-PCR) analysis of *ILK5* and *P2K1* transcripts in 10 day-old seedling or 2 week-old plants grown under normal conditions. Total RNA was isolated from 10-day-old plants (root, rosette leaf; R.L and whole plant) or 6 week-old plants (cauline leaf; C.L, stem and flower) and 2 μg of total RNA were used for qRT-PCR. The *SAND* reference gene was used for data normalization. Each value represents the means ± SD (n = 4). All above experiments were repeated two times with similar results. (*E*) qRT-PCR analysis of *ILK5* and *P2K1* gene expression in response to various abiotic stress conditions. Data shown as mean ± SEM, n = 2, **** *p*<0.0001, *** *p*<0.001, ** *p*<0.01, * *p*<0.05, *p*-value indicates significance relative to mock treatment of wild-type plants and was determined and analyzed using the GraphPad Prism 8 by one-way ANOVA followed by Dunnett’s multiple comparisons.. The *SAND* reference gene was used for data normalization. All above experiments were repeated two times with similar results.

**Fig. S7.**
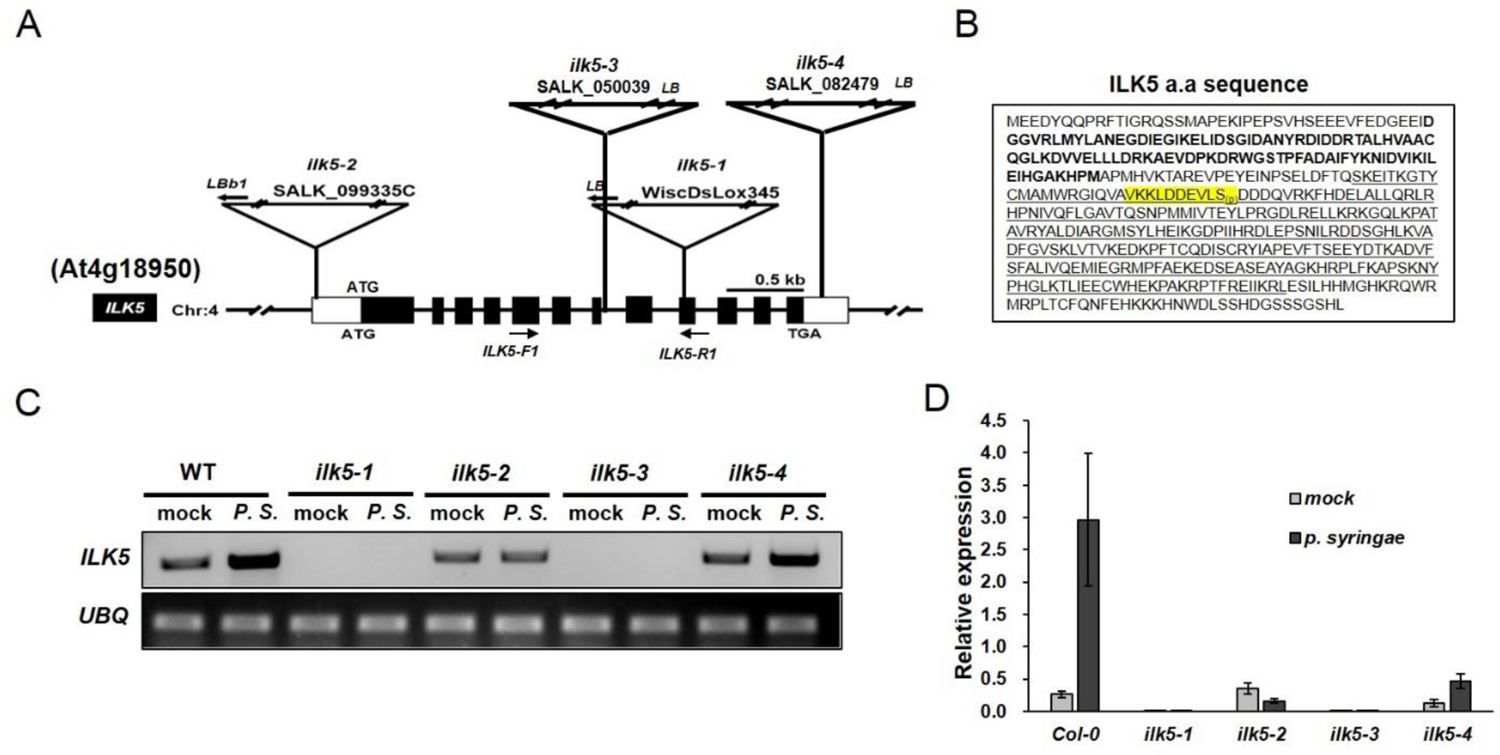
Schematic structure and sequence of *ILK5*. (*A*) Schematic diagram of *ILK5* (At4g18950) gene and T-DNA insertion regions. The filled boxes and the horizontal line between them represent exons and introns, respectively. The white boxes at both ends represent 5’ and 3’ UTRs. T-DNAs are inserted in the 9^th^ exon (*ilk5-1*; WiscDsLox345-348B17), 5’-UTR (*ilk5-2*; SALK_099335C), 7^th^ Intron (*ilk5-3;* SALK_050039) and 3’-UTR (*ilk5-4;* SALK_082479). (B) Protein sequence of *ILK5*. In the box, bold letter and underline indicate ANK and Ser/Thr kinase domains, respectively. Yellow highlighted region indicates phospho-peptide identified in the KiC assay. (*C* and *D*) RT-PCR (panel *C*) and real-time qRT-PCR (panel *D*) analysis of *ilk5-1*, *ilk5-2, ilk5-3* and *ilk5-4* mutants under *p. syringae* treatment. Location of used primer pairs is shown in panel *A*. qRT-PCR analysis of 3 week-old *ilk5* mutant plants. The *UBQ* or *SAND* reference gene was used for data normalization.

**Fig. S8.**
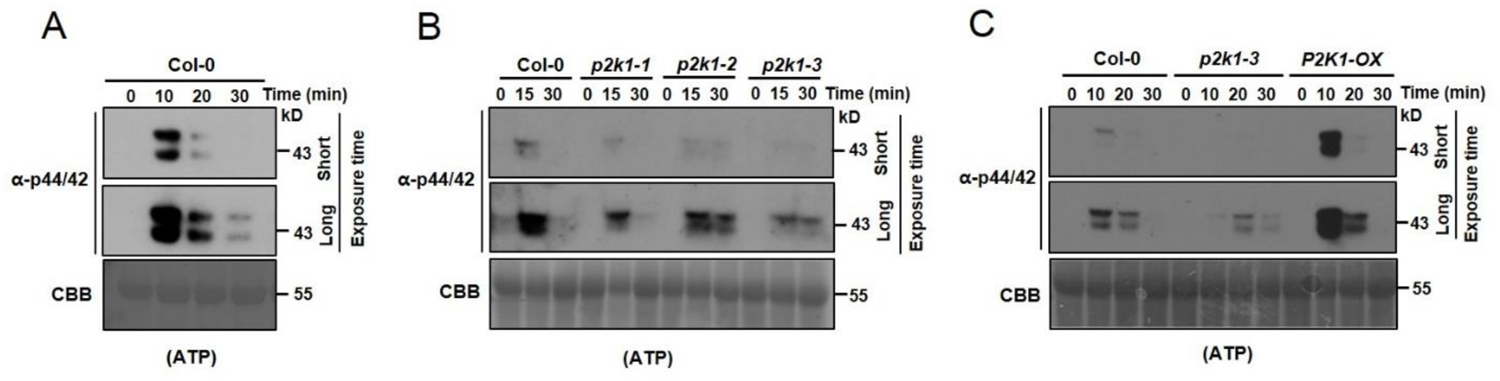
MPK3/6 phosphorylation is regulated by P2K1 under ATP treatment. (*A*) Phosphorylation of MPK3/6 was detected by western blotting using anti-phospho 44/42 antibody after treatment with ATP over a time course. CBB staining of protein was used as a loading control. (*B* and *C*) Phosphorylation of MPK3/6 were detected in two EMS mutagenized *p2k1* mutants (*p2k1-1*; D572N, *p2k1-2*; D525N) and a T-DNA insertion mutant (*p2k1-3*; Salk_042209), as well as a transgenic line ectopically, over-expressing P2K1. An anti-phospho 44/42 antibody was used to detect MPK3/6 phosphorylation. CBB staining of protein was used as a loading control.

**Fig. S9.**
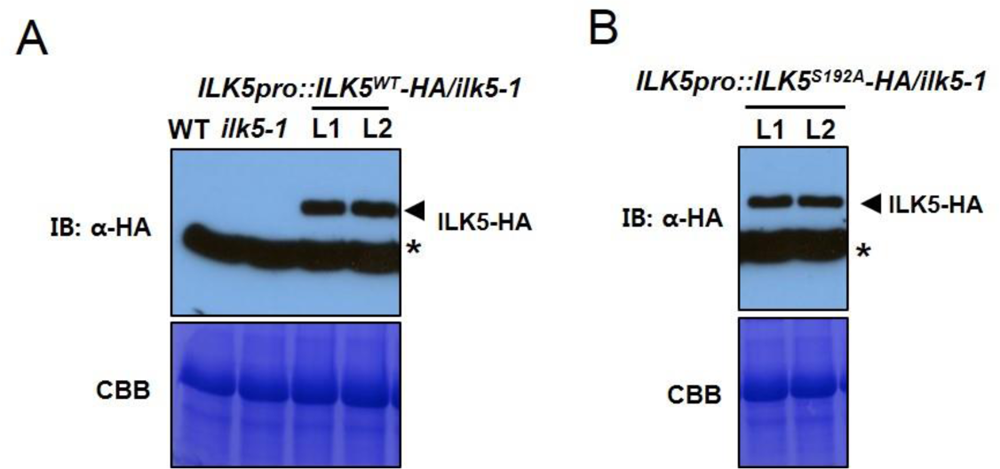
ILK5^WT^ and ILK5^S192A^ protein expression was determined by western blotting using anti-HA antibody. (*A* and *B*) Total protein was extracted from complemented lines in which the ILK5^WT^ or ILK5^S192A^ protein were expressed from the *ILK5* native promoter (L1 and L2). CBB staining of protein was used as a loading control. Asterisk represents non-specific bands.

**Fig. S10.**
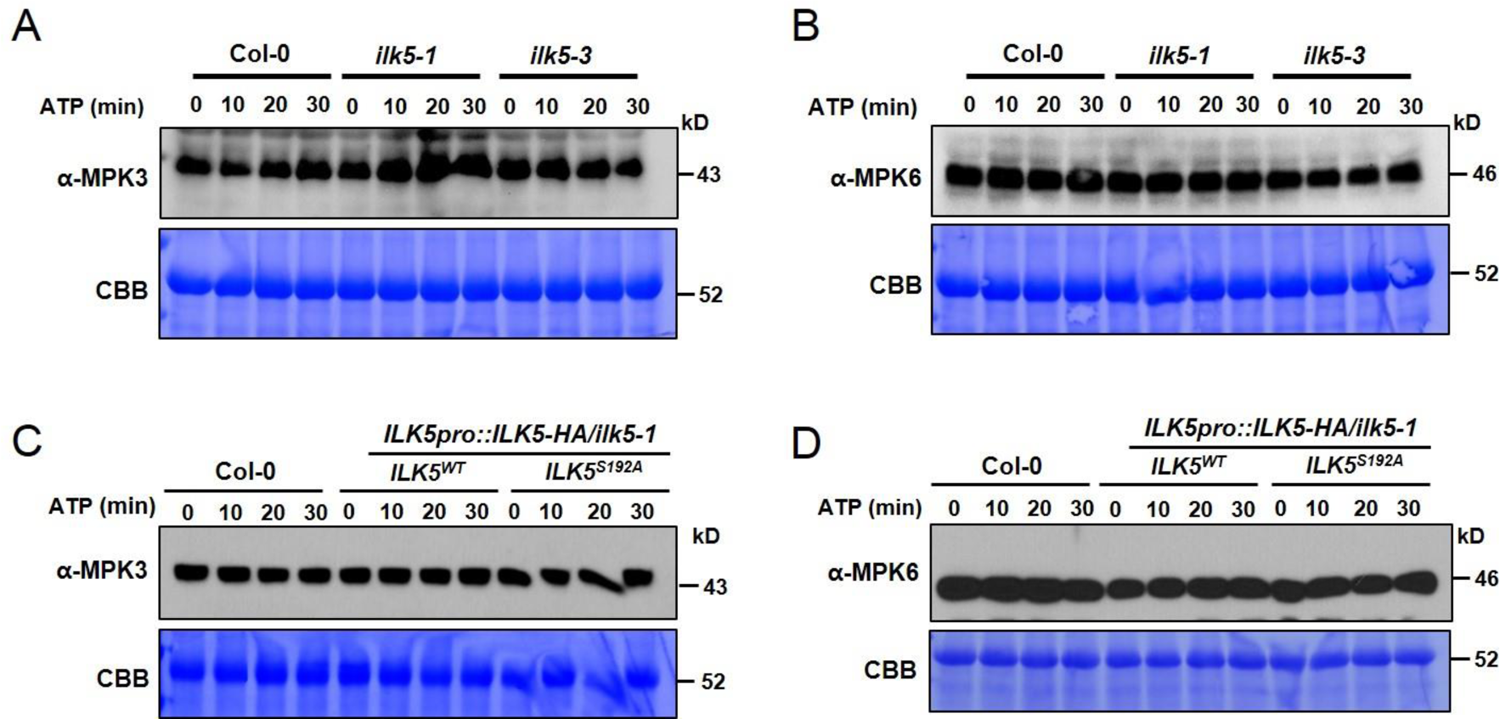
Investigation of MPK3/6 protein stability upon ATP treatment. (*A*-*D*) MPK3 (in panel *A* and *C*) and MPK6 (in panel *B* and *D*) was detected by western blotting using anti-MPK3 or -MPK6 antibodies after treatment with ATP over time in *ilk5-1* or *ilk5-3* mutants (panel *A* and *B*) or ILK5^WT^-HA or ILK5^S192A^-HA complemented lines (panel *C* and *D*). CBB staining of protein was used as a loading control.

**Fig. S11.**
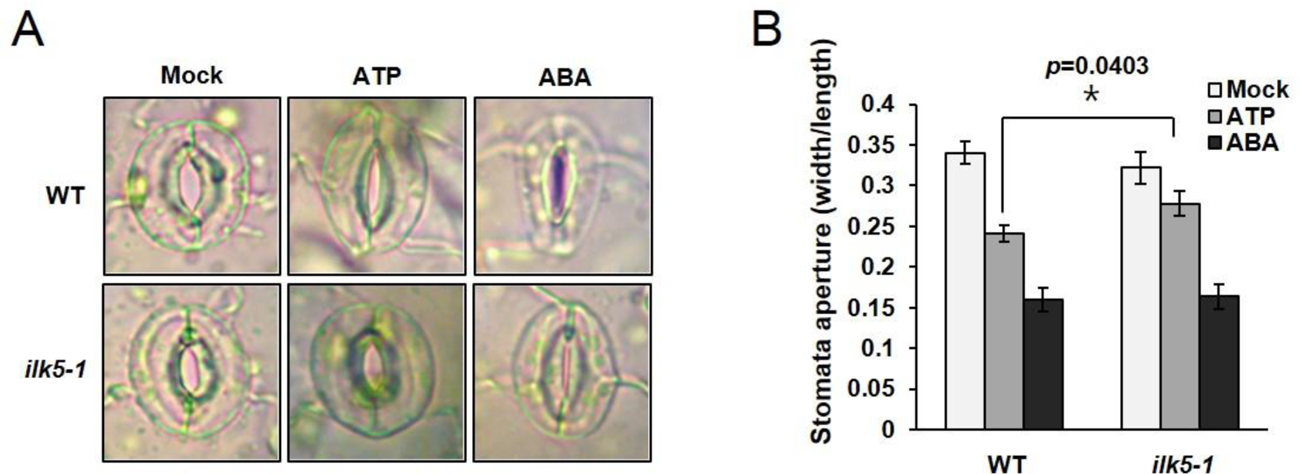
ILK5 is involved in stomatal closure in response to ATP treatment. (*A*, *B*) ILK5 is required for ATP-induced stomatal closure. Stomatal aperture was measured after treatment with 2 mM ATP or 5 μM ABA. Data shown as mean ± SEM, n ≥ 41, * *p*<0.05, *p*-value indicates significance relative to wild-type and was determined and analyzed using the GraphPad Prism 8 program by unpaired Student’s *t* test; two-way.

**Table S1.**
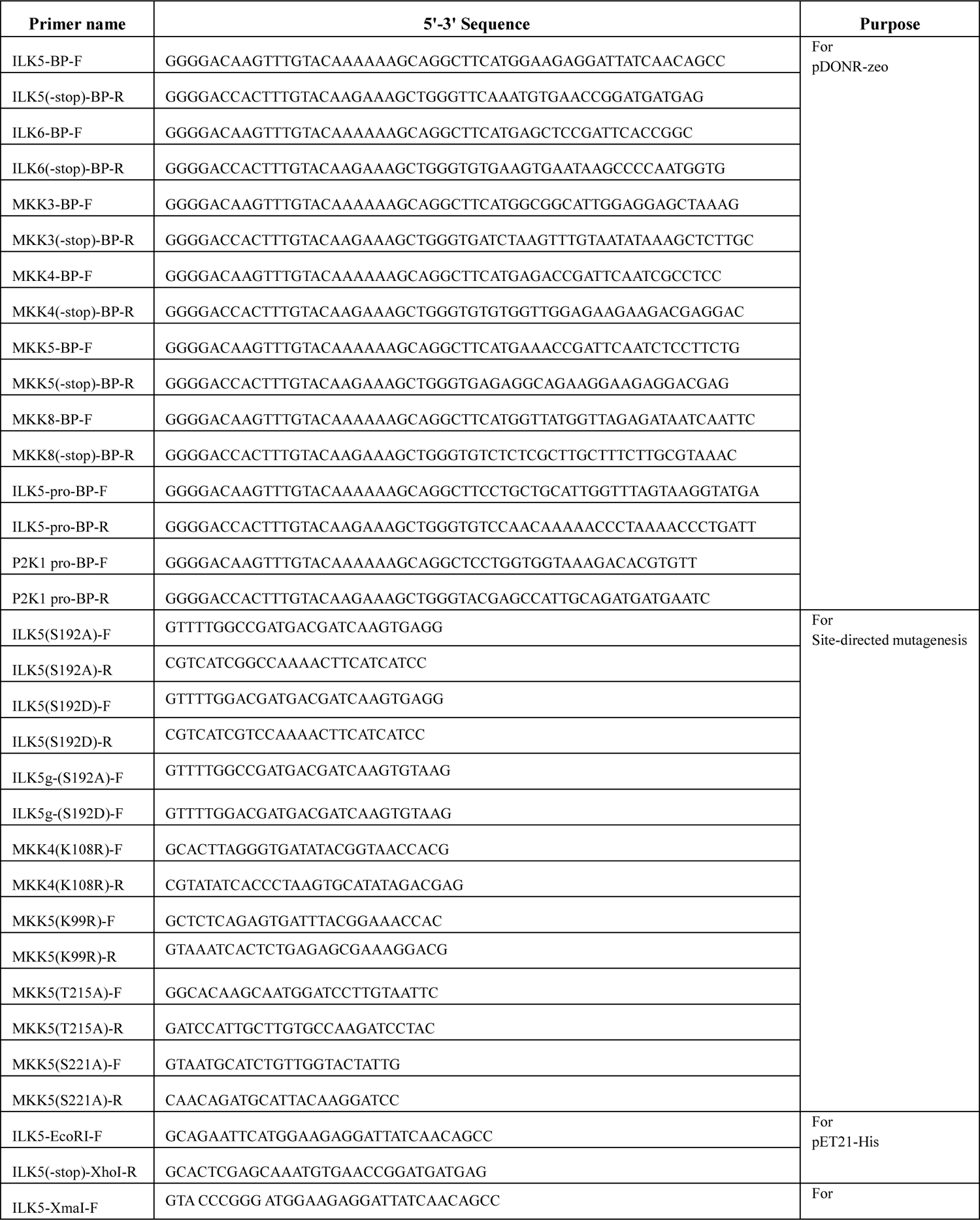
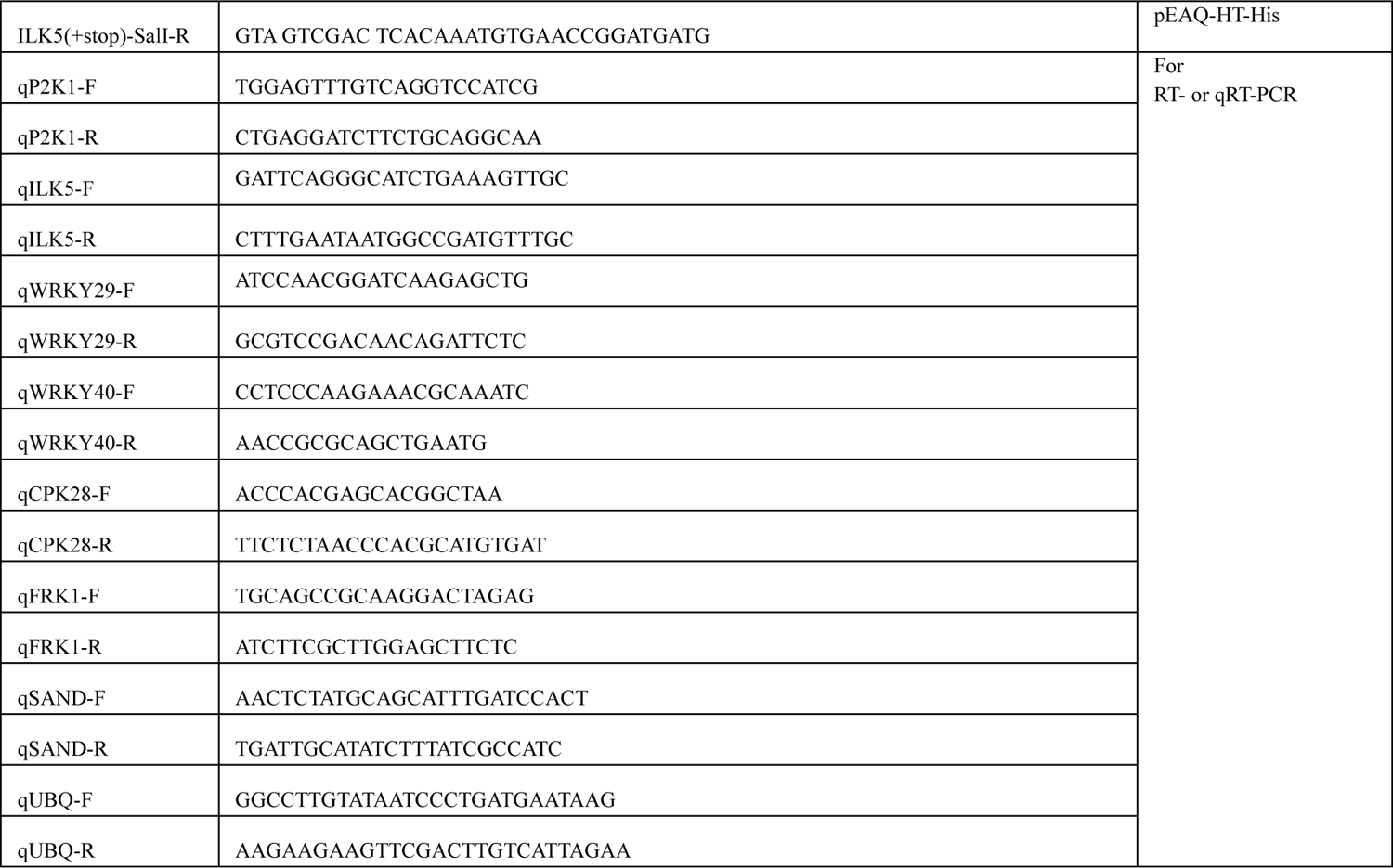
Sequence of primers used in this study

## Notes

### Competing Interest Statement

The authors have declared no competing interest.

## References

1. Y. Cao, K. Tanaka, C. T. Nguyen, G. Stacey, Extracellular ATP is a central signaling molecule in plant stress responses. Curr Opin Plant Biol 20, 82–87 (2014).

2. T.-T. T. Le, et al., Purinergic Signaling in Pulmonary Inflammation. Front. Immunol. 10 (2019).

3. R. Faria, L. Ferreira, R. Bezerra, V. Frutuoso, L. Alves, Action of natural products on p2 receptors: a reinvented era for drug discovery. Molecules 17, 13009–13025 (2012).

4. D. Ferrari, E. N. McNamee, M. Idzko, R. Gambari, H. K. Eltzschig, Purinergic Signaling During Immune Cell Trafficking. Trends Immunol 37, 399–411 (2016).

5. P. Matzinger, Friendly and dangerous signals: is the tissue in control? Nat Immunol 8, 11–13 (2007).

6. C. Cekic, J. Linden, Purinergic regulation of the immune system. Nat Rev Immunol 16, 177–192 (2016).

7. C. J. Song, I. Steinebrunner, X. Wang, S. C. Stout, S. J. Roux, Extracellular ATP induces the accumulation of superoxide via NADPH oxidases in Arabidopsis. Plant Physiol 140, 1222–1232 (2006).

8. D. Chen, et al., Extracellular ATP elicits DORN1-mediated RBOHD phosphorylation to regulate stomatal aperture. Nat Commun 8, 2265 (2017).

9. S. Deng, et al., Populus euphratica APYRASE2 Enhances Cold Tolerance by Modulating Vesicular Trafficking and Extracellular ATP in Arabidopsis Plants. Plant Physiol 169, 530–548 (2015).

10. C. Thomas, et al., A Role for Ectophosphatase in Xenobiotic Resistance. Plant Cell 12, 519–534 (2000).

11. W. Tang, S. R. Brady, Y. Sun, G. K. Muday, S. J. Roux, Extracellular ATP inhibits root gravitropism at concentrations that inhibit polar auxin transport. Plant Physiol 131, 147–154 (2003).

12. S.-Y. Kim, M. Sivaguru, G. Stacey, Extracellular ATP in plants. Visualization, localization, and analysis of physiological significance in growth and signaling. Plant Physiol 142, 984–992 (2006).

13. R. R. Lew, J. D. W. Dearnaley, Extracellular nucleotide effects on the electrical properties of growing Arabidopsis thaliana root hairs. Plant Science 153, 1–6 (2000).

14. R. Zhu, et al., Heterotrimeric G Protein-Regulated Ca2+ Influx and PIN2 Asymmetric Distribution Are Involved in Arabidopsis thaliana Roots’ Avoidance Response to Extracellular ATP. Front Plant Sci 8, 1522 (2017).

15. R. R. Weerasinghe, et al., Touch induces ATP release in Arabidopsis roots that is modulated by the heterotrimeric G-protein complex. FEBS Lett 583, 2521–2526 (2009).

16. S. Chivasa, B. K. Ndimba, W. J. Simon, K. Lindsey, A. R. Slabas, Extracellular ATP functions as an endogenous external metabolite regulating plant cell viability. Plant Cell 17, 3019–3034 (2005).

17. H. Feng, D. Guan, J. Bai, K. Sun, L. Jia, Extracellular ATP: a potential regulator of plant cell death. Mol Plant Pathol 16, 633–639 (2015).

18. J. Choi, et al., Identification of a Plant Receptor for Extracellular ATP. Science 343, 290–294 (2014).

19. J. Choi, et al., Extracellular ATP, a danger signal, is recognized by DORN1 in Arabidopsis. Biochem J 463, 429– 437 (2014).

20. C. T. Nguyen, et al., Computational Analysis of the Ligand Binding Site of the Extracellular ATP Receptor, DORN1. PLOS ONE 11, e0161894 (2016).

21. K. Bouwmeester, et al., The lectin receptor kinase LecRK-I.9 is a novel Phytophthora resistance component and a potential host target for a RXLR effector. PLoS Pathog 7, e1001327 (2011).

22. C. Balagué, et al., The Arabidopsis thaliana lectin receptor kinase LecRK-I.9 is required for full resistance to Pseudomonas syringae and affects jasmonate signalling. Mol Plant Pathol 18, 937–948 (2017).

23. K. Bouwmeester, et al., The Arabidopsis lectin receptor kinase LecRK-I.9 enhances resistance to Phytophthora infestans in Solanaceous plants. Plant Biotechnol J 12, 10–16 (2014).

24. Y. Wang, D. L. Nsibo, H. M. Juhar, F. Govers, K. Bouwmeester, Ectopic expression of Arabidopsis L-type lectin receptor kinase genes LecRK-I.9 and LecRK-IX.1 in Nicotiana benthamiana confers Phytophthora resistance. Plant Cell Rep 35, 845–855 (2016).

25. R. J. Myers, Y. Fichman, G. Stacey, R. Mittler, Extracellular ATP plays an important role in systemic wound response activation. 2022.01.06.475278 (2022).

26. D. Chen, et al., S-acylation of P2K1 mediates extracellular ATP-induced immune signaling in Arabidopsis. Nat Commun 12, 2750 (2021).

27. H. N. Duong, et al., Cyclic Nucleotide Gated Ion Channel 6 is involved in extracellular ATP signaling and plant immunity. Plant J (2021) https:/doi.org/10.1111/tpj.15636.

28. L. Wang, et al., Arabidopsis thaliana CYCLIC NUCLEOTIDE-GATED CHANNEL2 mediates extracellular ATP signal transduction in root epidermis. New Phytol (2022) https:/doi.org/10.1111/nph.17987.

29. A. Q. Pham, S.-H. Cho, C. T. Nguyen, G. Stacey, Arabidopsis Lectin Receptor Kinase P2K2 Is a Second Plant Receptor for Extracellular ATP and Contributes to Innate Immunity. Plant Physiol 183, 1364–1375 (2020).

30. K. Thulasi Devendrakumar, X. Li, Y. Zhang, MAP kinase signalling: interplays between plant PAMP- and effector-triggered immunity. Cell Mol Life Sci 75, 2981–2989 (2018).

31. M. CristinaRodriguez, M. Petersen, J. Mundy, Mitogen-Activated Protein Kinase Signaling in Plants. Annu Rev Plant Biol 61, 621–649 (2010).

32. MAPK Group, Mitogen-activated protein kinase cascades in plants: a new nomenclature. Trends Plant Sci 7, 301–308 (2002).

33. A. Champion, A. Picaud, Y. Henry, Reassessing the MAP3K and MAP4K relationships. Trends Plant Sci 9, 123–129 (2004).

34. J. Colcombet, H. Hirt, Arabidopsis MAPKs: a complex signalling network involved in multiple biological processes. Biochem J 413, 217–226 (2008).

35. C. A. Frye, R. W. Innes, An Arabidopsis mutant with enhanced resistance to powdery mildew. Plant Cell 10, 947–956 (1998).

36. C. A. Frye, D. Tang, R. W. Innes, Negative regulation of defense responses in plants by a conserved MAPKK kinase. Proc Natl Acad Sci U S A 98, 373–378 (2001).

37. K. L. Clark, P. B. Larsen, X. Wang, C. Chang, Association of the Arabidopsis CTR1 Raf-like kinase with the ETR1 and ERS ethylene receptors. Proc Natl Acad Sci U S A 95, 5401–5406 (1998).

38. S.-D. Yoo, Y.-H. Cho, G. Tena, Y. Xiong, J. Sheen, Dual control of nuclear EIN3 by bifurcate MAPK cascades in C2H4 signalling. Nature 451, 789–795 (2008).

39. J. J. Kieber, M. Rothenberg, G. Roman, K. A. Feldmann, J. R. Ecker, CTR1, a negative regulator of the ethylene response pathway in Arabidopsis, encodes a member of the raf family of protein kinases. Cell 72, 427–441 (1993).

40. S. I. Gibson, R. J. Laby, D. Kim, The sugar-insensitive1 (sis1) mutant of Arabidopsis is allelic to ctr1. Biochem Biophys Res Commun 280, 196–203 (2001).

41. D. Tang, K. M. Christiansen, R. W. Innes, Regulation of Plant Disease Resistance, Stress Responses, Cell Death, and Ethylene Signaling in Arabidopsis by the EDR1 Protein Kinase. Plant Physiol 138, 1018–1026 (2005).

42. S. C. Popescu, E. K. Brauer, G. Dimlioglu, G. V. Popescu, Insights into the Structure, Function, and Ion-Mediated Signaling Pathways Transduced by Plant Integrin-Linked Kinases. Front Plant Sci 8 (2017).

43. G. E. Hannigan, et al., Regulation of cell adhesion and anchorage-dependent growth by a new β1-integrin-linked protein kinase. Nature 379, 91–96 (1996).

44. Y. Tu, Y. Huang, Y. Zhang, Y. Hua, C. Wu, A New Focal Adhesion Protein That Interacts with Integrin-Linked Kinase and Regulates Cell Adhesion and Spreading. J Cell Biol 153, 585–598 (2001).

45. G. E. Hannigan, P. C. McDonald, M. P. Walsh, S. Dedhar, Integrin-linked kinase: Not so ‘pseudo’ after all. Oncogene 30, 4375–4385 (2011).

46. L. Dagnino, Integrin-linked kinase: a Scaffold protein unique among its ilk. J Cell Commun Signal 5, 81–83 (2011).

47. B. Meder, et al., PINCH Proteins Regulate Cardiac Contractility by Modulating Integrin-Linked Kinase-Protein Kinase B Signaling▿. Mol Cell Biol 31, 3424–3435 (2011).

48. D. Chinchilla, et al., A mutant ankyrin protein kinase from Medicago sativa affects Arabidopsis adventitious roots. Functional Plant Biol. 35, 92 (2008).

49. D. Chinchilla, et al., Ankyrin protein kinases: a novel type of plant kinase gene whose expression is induced by osmotic stress in alfalfa. Plant Mol Biol 51, 555–566 (2003).

50. T. Ceserani, A. Trofka, N. Gandotra, T. Nelson, VH1/BRL2 receptor-like kinase interacts with vascular-specific adaptor proteins VIT and VIK to influence leaf venation. Plant J 57, 1000–1014 (2009).

51. E. K. Brauer, et al., The Raf-like Kinase ILK1 and the High Affinity K+ Transporter HAK5 Are Required for Innate Immunity and Abiotic Stress Response1[OPEN]. Plant Physiol 171, 1470–1484 (2016).

52. M. Hayashi, S.-I. Inoue, Y. Ueno, T. Kinoshita, A Raf-like protein kinase BHP mediates blue light-dependent stomatal opening. Sci Rep 7, 45586 (2017).

53. N. Ahsan, et al., A versatile mass spectrometry-based method to both identify kinase client-relationships and characterize signaling network topology. J Proteome Res 12, 937–948 (2013).

54. Y. Huang, et al., A quantitative mass spectrometry-based approach for identifying protein kinase clients and quantifying kinase activity. Anal Biochem 402, 69–76 (2010).

55. C. Wu, S. Dedhar, Integrin-linked kinase (ILK) and its interactors. J Cell Biol 155, 505–510 (2001).

56. K. Nemoto, et al., Autophosphorylation profiling of Arabidopsis protein kinases using the cell-free system. Phytochemistry 72, 1136–1144 (2011).

57. T. Asai, et al., MAP kinase signalling cascade in Arabidopsis innate immunity. Nature 415, 977–983 (2002).

58. S. C. Popescu, et al., MAPK target networks in Arabidopsis thaliana revealed using functional protein microarrays. Genes Dev 23, 80–92 (2009).

59. K. Yamada, et al., The Arabidopsis CERK1-associated kinase PBL27 connects chitin perception to MAPK activation. EMBO J 35, 2468–2483 (2016).

60. J. Choi, Identification of an Extracellular Adenosine 5’-triphosphate Receptor in Arabidopsis Thaliana (2013).

61. D. Tripathi, T. Zhang, A. J. Koo, G. Stacey, K. Tanaka, Extracellular ATP Acts on Jasmonate Signaling to Reinforce Plant Defense1[OPEN]. Plant Physiol 176, 511–523 (2018).

62. C. Nawrath, J.-P. Métraux, Salicylic Acid Induction–Deficient Mutants of Arabidopsis Express PR-2 and PR-5 and Accumulate High Levels of Camalexin after Pathogen Inoculation. The Plant Cell 11, 1393–1404 (1999).

63. J. T. Neary, et al., Mitogenic Signaling by ATP/P2Y Purinergic Receptors in Astrocytes: Involvement of a Calcium-Independent Protein Kinase C, Extracellular Signal-Regulated Protein Kinase Pathway Distinct from the Phosphatidylinositol-Specific Phospholipase C/Calcium Pathway. J Neurosci 19, 4211–4220 (1999).

64. G. Clark, et al., Extracellular Nucleotides and Apyrases Regulate Stomatal Aperture in Arabidopsis1[W][OA]. Plant Physiol 156, 1740–1753 (2011).

65. V. Salmaso, K. A. Jacobson, Purinergic Signaling: Impact of GPCR Structures on Rational Drug Design. ChemMedChem 15, 1958–1973 (2020).

66. R. O. Hynes, Integrins: Bidirectional, Allosteric Signaling Machines. Cell 110, 673–687 (2002).

67. J. S. Desgrosellier, D. A. Cheresh, Integrins in cancer: biological implications and therapeutic opportunities. Nat Rev Cancer 10, 9–22 (2010).

68. C. Kim, F. Ye, M. H. Ginsberg, Regulation of Integrin Activation. Annu Rev Cell Dev Biol 27, 321–345 (2011).

69. S. Bagchi, et al., The P2Y2 Nucleotide Receptor Interacts with αv Integrins to Activate Go and Induce Cell Migration. J. Biol. Chem. 280, 39050–39057 (2005).

70. A. P. Bye, J. M. Gibbins, M. P. Mahaut-Smith, Ca2+ waves coordinate purinergic receptor–evoked integrin activation and polarization. Sci. Signal. 13 (2020).

71. C. Knepper, E. A. Savory, B. Day, Arabidopsis NDR1 Is an Integrin-Like Protein with a Role in Fluid Loss and Plasma Membrane-Cell Wall Adhesion. Plant Physiol 156, 286–300 (2011).

72. S. Thevananther, et al., Extracellular ATP activates c-jun N-terminal kinase signaling and cell cycle progression in hepatocytes. Hepatology 39, 393–402 (2004).

73. T. Markou, G. Vassort, A. Lazou, Regulation of MAPK pathways in response to purinergic stimulation of adult rat cardiac myocytes. Mol Cell Biochem 242, 163–171 (2003).

74. A. Schultze-Mosgau, et al., Characterization of calcium-mobilizing, purinergic P2Y2 receptors in human ovarian cancer cells. Mol Hum Reprod 6, 435–442 (2000).

75. X. Meng, et al., A MAPK Cascade Downstream of ERECTA Receptor-Like Protein Kinase Regulates Arabidopsis Inflorescence Architecture by Promoting Localized Cell Proliferation. The Plant Cell 24, 4948– 4960 (2012).

76. T. Soyano, R. Nishihama, K. Morikiyo, M. Ishikawa, Y. Machida, NQK1/NtMEK1 is a MAPKK that acts in the NPK1 MAPKKK-mediated MAPK cascade and is required for plant cytokinesis. Genes Dev 17, 1055–1067 (2003).

77. G. Bi, et al., Receptor-Like Cytoplasmic Kinases Directly Link Diverse Pattern Recognition Receptors to the Activation of Mitogen-Activated Protein Kinase Cascades in Arabidopsis. The Plant Cell 30, 1543–1561 (2018).

78. X. Liang, J.-M. Zhou, Receptor-Like Cytoplasmic Kinases: Central Players in Plant Receptor Kinase–Mediated Signaling. Annu Rev Plant Biol 69, 267–299 (2018).

79. D. Lu, et al., A receptor-like cytoplasmic kinase, BIK1, associates with a flagellin receptor complex to initiate plant innate immunity. Proc Natl Acad Sci U S A 107, 496–501 (2010).

80. F. Soma, F. Takahashi, T. Suzuki, K. Shinozaki, K. Yamaguchi-Shinozaki, Plant Raf-like kinases regulate the mRNA population upstream of ABA-unresponsive SnRK2 kinases under drought stress. Nat Commun 11, 1373 (2020).

81. S. C. Popescu, E. K. Brauer, G. Dimlioglu, G. V. Popescu, Insights into the Structure, Function, and Ion-Mediated Signaling Pathways Transduced by Plant Integrin-Linked Kinases. Front Plant Sci 8, 376 (2017).

82. J. A. Adams, Kinetic and catalytic mechanisms of protein kinases. Chem Rev 101, 2271–2290 (2001).

83. J. Su, et al., Regulation of Stomatal Immunity by Interdependent Functions of a Pathogen-Responsive MPK3/MPK6 Cascade and Abscisic Acid. Plant Cell 29, 526–542 (2017).

## SI references

1. D. Chen, et al., Extracellular ATP elicits DORN1-mediated RBOHD phosphorylation to regulate stomatal aperture. Nat Commun 8, 2265 (2017).

2. S. J. Clough, A. F. Bent, Floral dip: a simplified method for Agrobacterium-mediated transformation of Arabidopsis thaliana. Plant J 16, 735–743 (1998).

3. D. W. Kim, S. J. Jeon, S. M. Hwang, J. C. Hong, J. D. Bahk, The C3H-type zinc finger protein GDS1/C3H42 is a nuclear-speckle-localized protein that is essential for normal growth and development in Arabidopsis. Plant Sci 250, 141–153 (2016).

4. Y. Cao, H. Li, A. Q. Pham, G. Stacey, An Improved Transient Expression System Using Arabidopsis Protoplasts. Curr Protoc Plant Biol 1, 285–291 (2016).

5. F. Sainsbury, E. C. Thuenemann, G. P. Lomonossoff, pEAQ: versatile expression vectors for easy and quick transient expression of heterologous proteins in plants. Plant Biotechnol J 7, 682–693 (2009).

6. A. de Souza, Expression and Partial Purification of His-tagged Proteins in a Plant System. Bio-protocol 5, e1572–e1572 (2015).

7. D. Lu, et al., A receptor-like cytoplasmic kinase, BIK1, associates with a flagellin receptor complex to initiate plant innate immunity. Proc Natl Acad Sci U S A 107, 496–501 (2010).

